# Commensal Bacteria in the Gut Microbiome Encode Previously Unrecognized Aromatic Hydrocarbon Catabolic Machinery

**DOI:** 10.64898/2026.10.09.758022

**Authors:** Chinenye P. Nwike, Oluwatosin P. Shittu, Madison E. Andrews, Evan C. Roos, Tulasi Rao Relangi, Aaron T. Wright

## Abstract

**Introduction:** Polycyclic aromatic hydrocarbon (PAH) toxicology has been primarily defined by host metabolic bioactivation to reactive intermediates. Yet ingested PAHs and biliary-delivered metabolites also enter the intestinal lumen, where they encounter the gut microbiota, an additional but understudied metabolic compartment that may alter toxicant fate. Although PAH interactions are well characterized in environmental microbes, resulting in defined enzymatic pathways for their catabolism, whether gut commensal organisms express related machinery in vivo remains poorly understood. To address this gap, we used an oral murine-exposure model of naphthalene and integrated compartment-resolved 16S rRNA gene profiling, global metaproteomics, and chemoproteomics to resolve the PAH-processing repertoire of the gut microbiota and its spatial distribution.

**Results:** Naphthalene, a common environmental toxicant and representative PAH, elicited a compartment-dependent functional response that was most pronounced in the large intestine. Distally, exposure was associated with 2,386 differential proteins and accounted for 48.4% of metaproteomic variation, compared with 7.9% of 16S rRNA gene compositional variation. The small intestine showed a reduced proteomic response, with 994 differentially abundant proteins. Hydrocarbon catabolic markers showed a 4.8-fold distal expansion (36 to 172 proteins), including the naphthalene-catabolic *nahE*, proteins associated with phenylacetate, β-ketoadipate, catechol/TOL, and anaerobic CoA-linked redox machinery. The small-intestinal repertoire decreased from 19 to 14 proteins, though aromatic-catabolic markers doubled, indicating a selective expansion. Critically, these homologs concentrated in Lachnospiraceae, a gut family that is prevalent in humans, primarily known for anaerobic fermentation, with no priorly described PAH-processing capacity. An uncultured *Acetatifactor* showed the largest hydrocarbon hits (14 to 82 protein increase), with additional contributions from *Kineothrix, Eisenbergiella,* and *Petralouisia*. Chemoproteomics, using a naphthalene-based probe, revealed enrichment of an orthogonal proteome of proteins interacting with naphthalene, capturing 171 small-intestinal and 67 large-intestinal proteins that were undetected in the global metaproteomes.

**Conclusions:** Gut commensals express homologs of aromatic-hydrocarbon catabolic enzymes characterized in environmental microorganisms, and naphthalene amplifies this repertoire most strongly in the large intestine. The decoupling of taxonomic structure, expressed-protein representation, and probe engagement identifies a functional dimension of PAH exposure that composition-based profiling does not resolve and positions microbiome variation as a candidate modifier of PAH metabolic fate.

## Introduction

Polycyclic aromatic hydrocarbons (PAHs) are a chemically diverse class of ubiquitous environmental pollutants whose biological effects are shaped by metabolic transformation. Humans encounter PAHs through inhalation, ingestion and dermal contact, and the individual compounds differ in environmental persistence, absorption and toxicological potency [1–5]. Prior mechanistic studies have thus characterized various mechanisms of host-mediated PAH metabolism and toxicity. Following uptake, host enzymes can convert them to diol epoxides, quinones and other reactive intermediates that drive oxidative stress, DNA damage, immune dysfunction, developmental toxicity and carcinogenesis [1, 5–11]. However, a poorly explored consequence of host PAH exposure is its delivery to the intestinal lumen via direct oral ingestion, swallowed mucociliary-cleared material and biliary delivery of host-derived metabolites and conjugates [12–15]. These routes place the gut microbiota at an interface where it encounters and interacts with these toxicants alongside host enzymes and represents an additional metabolic compartment that may alter PAH persistence, chemical form, absorption and toxicological fate during host exposure. Naphthalene is particularly relevant to this interface due to its prominence in human exposure profiles. It accounted for more than 60% of summed PAH concentrations in paired indoor and outdoor measurements [16], and its metabolite 2-hydroxynaphthalene was the most abundant urinary PAH marker across seven sites in three United States pregnancy cohorts [17]. Despite this exposure burden, the metabolic landscape of gut microbial interactions with such PAHs remains poorly resolved.

Existing studies have shown PAH exposure perturbs the intestinal microbiome and accompanies changes in intestinal physiology and clinical outcomes. In mice, oral benzo[a]pyrene exposure has altered gut community structure alongside ileal and colonic inflammatory changes [18]. Benzo[a]pyrene can also alter microbial transcription without extensive compositional restructuring [18, 19]. In humans, PAH burden has been associated with gastrointestinal disease prevalence and Crohn’s disease [3, 11]. Evidence for microbial PAH transformation is more limited. Ex vivo analyses have revealed that human colon microbiota can transform parent PAHs, including naphthalene, into estrogenic products [20]. Separately, a metagenomic analysis of pollution-exposed human populations identified enrichment of genes associated with xenobiotic degradation and resistance [21]. Together, the existing evidence of gut microbial PAH interactions has remained predominantly descriptive and conspicuously disconnected from the biochemical architecture underlying the interactions. This underscores the need for protein-level resolution to functionally interrogate the expressed systems, metabolic representation, and taxa associated with PAH processing in the gut.

Biochemical precedent for microbial PAH metabolism has been established in environmental systems, such as sediment and soil [22]. Under aerobic conditions, bacteria use ring-hydroxylating and ring-cleavage enzymes to catabolize and route aromatic-derived carbon into central metabolism [22–25]. Anaerobic naphthalene-metabolizing microorganisms use distinct mechanisms that include carboxylation, formation and reductive dearomatization of 2-naphthoyl-CoA, and subsequent CoA-transferase, acyl-CoA dehydrogenase and β-oxidation-like reactions [26–29]. However, whether similar or related metabolic machinery is expressed by native mammalian gut commensals remains largely unknown. Oxygen availability, substrate supply, microbial biomass and transit time vary along the gastrointestinal tract [30]. The small intestine contains a comparatively oxygenated tissue interface, rapid transit and high exposure to host secretions, whereas the large intestine supports a dense, predominantly anaerobic community with prolonged substrate residence and extensive fermentation [30, 31]. These spatial differences can influence which and how organisms and proteins interact with PAHs or their host metabolites in the intestinal lumen and thus provide a strong rationale for a compartment-resolved interrogation.

To functionally annotate the metabolic capacity of commensal gut bacteria toward PAH exposure, we combined compartment-resolved 16S rRNA gene profiling, global metaproteomics, and chemoproteomics to mice orally exposed to naphthalene (**Fig. 1**). Building on our laboratory’s resin-immobilized probe platform for complex microbial proteomes [32], we also developed naphthalene-derived probes to enrich PAH-interacting enzymes from the gut proteome. With these, we interrogated whether naphthalene exposure produces functional changes not predicted by community composition, whether gut commensals express proteins associated with aromatic-hydrocarbon processing, and how this repertoire is distributed across intestinal compartments and taxa. We hypothesized that proteomic profiling would uncover metabolic adaptations for aromatic processing in the gut, revealing functional capabilities which may be invisible to compositional analyses.

**Figure 1.**
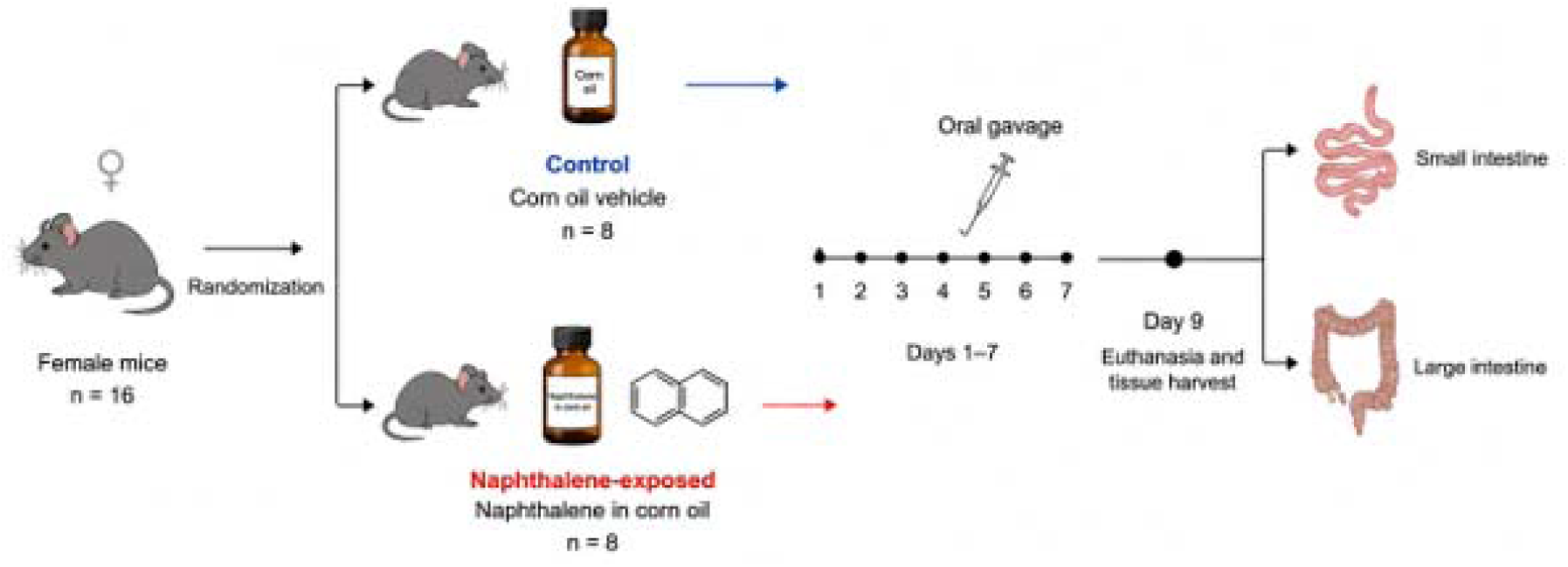
Experimental design for acute oral naphthalene exposure to mice. Mice were split into two groups of eight (8) mice each and oral gavaged with corn oil or naphthalene in corn oil for seven days. All mice were euthanized on day nine (9) and intestinal contents were harvested.

## Methods

### Materials and Reagents

All chemicals were purchased and used without further purification. Non-pharmaceutical grade naphthalene, TentaGel-XV-NH_2_ (0.27 mmol/g), and 10 mL Torviq reaction vessel for solid phase organic syntheses were purchased from Fisher Scientific. Corn oil was purchased from Sigma Aldrich. 10 mL fritted peptide vessel was purchased from Chemglass. PEG spacer (Fmoc-N-amido-PEG_4_-OH) was purchased from BroadPharm. 6-Bromo-2-Naphthoic acid was purchased from Ambeed. All other solvents and reagents were purchased from Fisher Scientific, Avantor, and Sigma Aldrich.

### Probe Synthesis

Details for probe synthesis and characterization data are in the Supplementary Information.

### Mice Study Design

All animal procedures were conducted based on an approved Institutional Animal Care and Use Committee (IACUC) protocol #2355412-2 at Baylor University. Sixteen seven-week-old female C57BL/6J mice were purchased from The Jackson Laboratory and housed in the vivarium at Baylor University. Mice had access to standard mice chow and water *ad libitum*. Mice were acclimatized for a one-week period before being randomly split into different cages for naphthalene-treatment or control groups of eight mice. The treatment group received a daily oral gavage of naphthalene (0.20 mL) in corn oil at 180 μmol/kg body weight for one-week. The control group received an equal volume of corn oil on the same dosing schedule.

### Sample Collection

Following the one-week dosing period, mice were kept one day without treatment prior to euthanasia. Mice were anesthetized under isoflurane exposure before being euthanized via cervical dislocation. Post-euthanasia, large and small intestines of the naphthalene-treated and control mice were immediately extracted under sterile and ice-cold conditions. Gut intestinal samples were split into tubes for downstream 16S sequencing, metaproteomics and chemoproteomics processing. Sample tubes were flash-frozen using liquid nitrogen and stored in −80 °C until further processing.

### 16S rRNA Amplicon Long-read Sequencing of Gut Microbes

To resolve microbial community structure at high taxonomic resolution, extraction of microbial DNA from the murine small and large intestinal contents (n = 8 replicates per naphthalene exposed or control group) was performed using the ZymoBIOMICS 16S rRNA Long Read Library Prep Kit (Zymo Research), following the manufacturer’s protocol. The extracted DNA was normalized to 1 ng/μL in nuclease-free water and used as input for full-length 16S rRNA gene amplification and barcoded library construction with the Quick-16S Full-Length Library Prep Kit, Primer Set 1 (D6450-PS1, Zymo Research), according to the manufacturer’s instructions. The purified library was converted into nanopore-compatible libraries using an Oxford Nanopore Technologies (ONT) Ligation Sequencing Kit. The protocol involved end-pair and dA-tailing of the double-stranded DNA fragments, followed by the ligation of sequencing adapters pre-loaded with motor proteins required for controlled pore translocation rate. The final ligated library (50 ng) was loaded onto an R10.4.1 MiniON flow cell and sequenced on a MiniON Mk1B (ONT) device.

Basecalling was disabled during sequencing because libraries were prepared with custom barcodes and primers from the Quick-16S Full-Length Library Prep Kit. Post-sequencing, raw signal data were generated in POD5 format and basecalled post-run with Dorado (dna_r10.4.1_e8.2_400bps_hac@v5.2.0; ONT) to produce FASTQ files. Reads were demultiplexed according to the manufacturer’s recommendations using the MiniBar Python script (minibar.py, downloaded from https://github.com/calacademy-research/minibar/blob/master/minibar.py), which performs dual barcode and primer-based demultiplexing of nanopore amplicon reads. Resulting FASTQ files were processed with an *in-house* shell script to trim the barcodes and retain only reads with mean quality score ≥ 9 and sequence lengths between 950 and 1,750 bp. to emit FASTQ files.

Assigning taxonomy and abundance estimation of filtered FASTQ files were done with EMU (version 3.5.1)[33], using an *in-house* custom EMU database that was curated from 16s rRNA full length sequence databases including SILVA 138.2 release[34], GTDB 226 release[35], and MIMt M2c_25_10 release[36]. Output files, including the taxonomically assigned feature abundance table generated by EMU and corresponding sample metadata, were imported into R and integrated using the phyloseq package for downstream analyses. Alpha diversity metrics (Shannon diversity, observed richness and Simpson diversity), were computed, and group differences were assessed using the non-parametric Wilcoxon rank-sum test. Beta-diversity was estimated using Bray Curtis dissimilarity distances, and overall community differences were evaluated via PERMANOVA. Sample clustering and separation were then visualized using a principal coordinates analysis (PCoA) plot. Microbial composition was summarized using relative abundance bar plots at the family, genus, and species level generated with ggplot2. Differential abundance analysis was performed using ANCOM-BC2 on 16S rRNA-derived count data.

### Microbial Protein Extraction and Proteomic Sample Preparation

Detailed procedures for bacterial cultivation, microbial protein extraction, proteolytic digestion, resin-based enrichment and peptide preparation are provided in the Supplementary Information. Briefly, *Pseudomonas putida* G7 was used as an induced reference for naphthalene-catabolic proteins. Overnight cultures grown in glucose-tryptone-yeast extract medium were diluted 1:75 into naphthalene–tryptone– yeast extract medium and cultured at 28 °C to mid-logarithmic phase (OD600 ≈ 0.7). Cells were harvested, washed in phosphate-buffered saline (PBS), disrupted by bead beating and clarified by centrifugation.

For murine metaproteomics, small-and large-intestinal contents were homogenized in ice-cold PBS and clarified by low-speed centrifugation. The resulting microbial-enriched fractions were disrupted by bead beating and clarified by centrifugation. Protein concentrations in the bacterial and intestinal lysates were determined using a bicinchoninic acid (BCA) assay.

For global proteomic analysis, 100 µg protein from each murine sample (n = 8 per exposure/control group and intestinal segment) was denatured in 8 M urea, reduced with dithiothreitol, alkylated with iodoacetamide and digested overnight with sequencing-grade trypsin at a 1:50 enzyme-to-protein ratio. Peptides were desalted by reversed-phase C18 solid-phase extraction, dried, resuspended in 0.1% formic acid containing 2% acetonitrile, quantified fluorometrically and normalized before Liquid Chromatography-Tandem Mass Spectrometry (LC-MS/MS) analysis.

#### Resin-based Chemoproteomic Enrichment

Microbial lysates were normalized to 500 µg total protein and incubated for 90 min at 37 °C with the applicable naphthalene-derived resin probe, **Br-NAP** or **DADPS-Br-NAP**, or with the acetyl-capped control resin, **CR-C** (**Scheme S1,S2; Fig. 6a–c**). Resin-bound proteins were denatured in SDS and subjected to sequential stringent washes to remove unbound and nonspecifically associated proteins. Enriched proteins were reduced with tris(2-carboxyethyl)phosphine (TCEP), alkylated with iodoacetamide and digested on-bead overnight with trypsin. Released peptides were collected, dried, resuspended in 0.1% formic acid containing 2% acetonitrile, clarified by ultracentrifugation and transferred to autosampler vials for LC-MS/MS analysis. Detailed procedures for peptide separation and acquisition are provided in the Supplementary Information.

### Protein Identification and Bioinformatics Data Processing

Raw MS/MS spectra were searched using an in-house MS-GF+ search engine pipeline (https://github.com/thulasis/MSGFPlus_Pipeline) which includes raw file conversion using ThermoRawFileParser v1.4.4 [37] and intensity assignments against curated protein sequence databases tailored to each experiment [38, 39]. For murine intestinal metaproteomes, spectra were searched against an *in-house* FASTA database constructed from reference proteomes of taxa identified via 16S rRNA gene profiling, supplemented with the Global Proteome Machine “common Repository of Adventitious Proteins” (cRAP) contaminant FASTA, which lists the common contaminant proteins observed in proteomics experiments[40]. For *P. putida* G7 samples, spectra were queried against the *P. putida* G7 reference proteome assembled from UniProt and similarly augmented with contaminants. A target–decoy strategy was employed during database construction to enable downstream score calibration and validation.

Search parameters included trypsin specificity with up to two missed cleavages, carbamidomethylation of cysteine as a fixed modification, and methionine oxidation and N-terminal acetylation as variable modifications. Peptide-spectrum matches (PSMs) were filtered based on MSMSScore, defined as the negative logarithm of the spectral E-value (−log10 SpecEValue), to retain high-confidence identifications. Peptides were subsequently aggregated to proteins using the Reference Rollup (RRollup) method implemented within the MS-GF+ pipeline. Protein identifications were retained if supported by at least one unique peptide and an MSMSScore ≥ 8 [38].

### Quantitative Metaproteomics and Chemoproteomics Data Analyses

#### Protein-intensity processing

Metaproteomic analyses were performed independently for the small and large intestine using eight mice per exposure/control group. Murine chemoproteomic analyses included four biological replicates per probe-by-exposure group, and *P. putida* G7 chemoproteomic analyses included four independent cultures per resin group. Proteins were retained when quantified in at least half of the replicates of one experimental group. Zero intensities were treated as missing values (see Supplementary Information), and retained intensities were log2 transformed. Samples were median-centered within their respective experimental groups.

Two versions of each intensity matrix were retained. The normalized non-imputed matrix, denoted NI, retained missing observations and was used for detection, presence and absence classification, abundance aggregation, protein-to-KO summation, and taxonomic abundance summaries. The imputed matrix, denoted IM, was used for analyses requiring a complete feature matrix, including protein-level linear modelling, principal component analysis, hierarchical clustering and row-standardized heatmaps.

#### Differential protein-abundance analysis

Differential protein abundance was analyzed in limma v3.66.0 [41]. Moderated statistics were calculated using empirical Bayes variance estimation with intensity-dependent trend modelling and robust estimation. *P* values were adjusted separately for each prespecified contrast using the Benjamini–Hochberg procedure.

#### Metaproteomic analysis

For the murine metaproteomes, naphthalene-exposed and control mice were compared separately within each intestinal segment. Proteins were considered differentially abundant at an adjusted *P* value < 0.05 and an absolute log2 fold change ≥ 2. Positive fold changes indicate greater abundance in exposed sample groups. Proteins detected in at least four replicates of one condition and in none of the other were additionally classified as condition-unique using the non-imputed data.

#### Murine chemoproteomic analysis

Murine chemoproteomic intensities were modeled using four groups defined by sample group and resin treatment: **Br-NAP** and **CR-C** in naphthalene-exposed mice and **Br-NAP** and **CR-C** in unexposed control mice.

Two prespecified limma contrasts were tested. The first compared **Br-NAP** with **CR-C** in naphthalene-exposed animals, defined as [Br-NAP_exposed − CR-C_exposed]. The second compared **Br-NAP** with **CR-C** in unexposed control mice, defined as [Br-NAP_control − CR-C_control]. Positive log2 fold changes therefore indicate preferential protein recovery with **Br-NAP** relative to **CR-C** within the corresponding sample group. Proteins were considered **Br-NAP**-enriched at an adjusted *P* value < 0.05 and a log2 fold change ≥ 1. Proteins meeting these criteria in exposed sample group were classified as “Probe Exposed”, those meeting the criteria in control group as “Probe Ctrl”. Proteins meeting the criteria in neither comparison were classified as not significantly probe-enriched. For descriptive comparison of enrichment magnitude between sample groups, the **Br-NAP**-versus-**CR-C** log2 fold change in control group was subtracted from that in exposed group. This difference was used to visualize the direction and magnitude of exposure-associated changes in probe recovery.

#### Functional annotation and KO-level analysis

Detailed description of KEGG Orthology (KO) assignment criteria and analyses is included in the Supplementary Information. In brief, identifiers were assigned independently using KofamScan v1.3.0 and eggNOG-mapper v2.1 [42, 43]. For each protein, KO assignments were prioritized as follows: KofamScan hits meeting the model-specific adaptive threshold, eggNOG-mapper assignments, and KofamScan hits reaching at least 50% of the model-specific threshold. KO names, enzyme commission numbers, BRITE hierarchies and pathway memberships were retrieved from KEGG on 07-13-2026.

#### Hydrocarbon-degradation annotation

Hydrocarbon-degradation homologues were annotated using CANT-HYD profile hidden Markov model collection (37 models; repository accessed 3rd March 2026) and the November 2023 release of the Hydrocarbon Aerobic Degradation Enzymes and Genes (HADEG) database. CANT-HYD profile hidden Markov models were searched using HMMER hmmsearch v3.4 and default reporting thresholds. HADEG sequences were searched using NCBI BLASTP v2.12.0+, and an E value threshold of 1 × 10^−5^. The highest-scoring match per protein was retained during downstream hit processing. The final applied score, coverage and curation criteria are provided in the deposited scripts and annotation tables. Complete hit-level results, including reference identifiers, scores, E values and assigned degradation categories, are provided in Supplementary Files 3 and 4.

## Results

### Protein architecture linked to naphthalene metabolism

The large intestine showed the dominant protein-level response to naphthalene despite showing the weakest compositional shift (**Supplementary Fig. 1,2**). Exposure explained 48.4% of large-intestinal metaproteomic variation but only 7.9% of the corresponding Bray-Curtis compositional variation (**Supplementary Fig. 2**). In the small intestine, the difference was narrower, with exposure explaining 45.6% of proteomic variation and 31.7% of compositional variation. The strongest separation between composition and protein representation therefore occurred distally.

Notably, exposure expanded the intestinal metaproteome while preserving a large fraction of the shared protein core. Shared proteins remained the largest fraction in both compartments, accounting for 1,542 of 2,074 proteins in the small intestine and 2,814 of 4,414 proteins in the large intestine (74.3% and 63.8%, respectively; **Fig. 2a,b**). However, exposure added 322 condition-exclusive proteins in the small intestine and 1,060 in the large intestine, compared with 210 and 540 proteins detected only in the corresponding controls. The exposure-exclusive fraction was therefore substantially larger distally, where it comprised 24.0% of all detected proteins, compared with 15.5% proximally.

**Figure 2.**
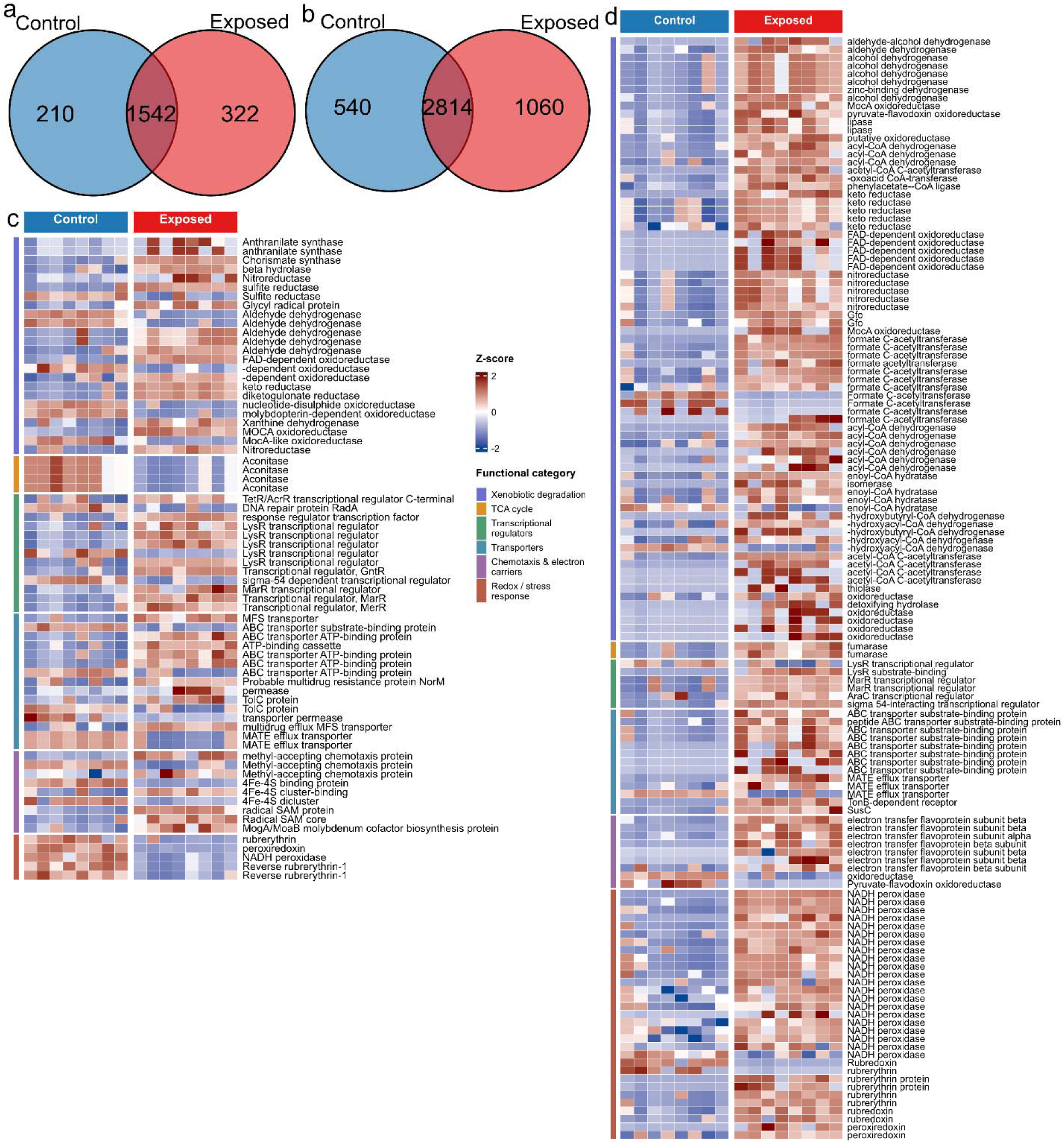
The intestinal metaproteome encodes compartment-specific aromatic-hydrocarbon catabolic potential. (**a,b**) Overlap of proteins detected in control and naphthalene-exposed samples from the small intestine (**a**) and large intestine (**b**). Values indicate proteins detected only in controls, in both conditions, or only following exposure. (**c,d**) Abundance profiles of selected candidate aromatic-catabolic proteins and associated metabolic, regulatory, transport and redox functions identified by LC-MS metaproteomics in the small intestine (**c**) and large intestine (**d**). Normalized protein intensities were standardized within each protein across control and exposed samples. Red and blue indicate values above and below the protein-specific mean, respectively. Each column represents one biological replicate, with eight mice per condition.

To determine whether the exposure-associated metaproteome encoded a protein architecture relevant to PAH transformation, we curated catabolic and supporting systems recurrently implicated in environmental microbial degradation of naphthalene and related aromatic xenobiotics. These comprised aromatic-compound transformation enzymes together with central carbon metabolism, transcriptional regulation, transport, chemotaxis, electron transfer and redox or stress-defense proteins. Environmental transcriptomic and proteomic studies indicate that these systems are differentially represented during aromatic degradation, although their direction and magnitude vary with organism, substrate and exposure conditions [44–47]. Proteins were grouped into broad categories: xenobiotic degradation, TCA-cycle metabolism, transcriptional regulation, transport, chemotaxis and electron transfer, and redox or stress defense (**Fig. 2c,d**). The large-intestinal profile organized around a coordinated CoA-linked catabolic and redox-supporting protein complement (**Fig. 2d**). Most selected xenobiotic-associated proteins increased in exposed mice, including acyl-CoA dehydrogenases, enoyl-CoA hydratases, hydroxyacyl-CoA dehydrogenases, acetyl-CoA acetyltransferases, CoA transferases and aldehyde dehydrogenases. Exposure-associated electron-transfer flavoprotein subunits co-occurred with the acyl-CoA dehydrogenases, consistent with transfer of reducing equivalents from FAD-dependent acyl-CoA oxidation into downstream electron-transfer pathways under distal anaerobic conditions [48, 49]. Rubrerythrins, rubredoxin and NADH peroxidases formed the dominant redox-defense complement while formate C-acetyltransferase and pyruvate-flavodoxin oxidoreductase further connected this profile to anaerobic carbon metabolism and low-potential electron transfer.

The small-intestinal profile yielded a different metabolic response to naphthalene exposure (**Fig. 2c**). Aldehyde dehydrogenases, nitroreductases, sulfite reductases and FAD-dependent oxidoreductases predominated among the selected xenobiotic-associated proteins, indicating localized oxidoreductive transformation and reactive-carbonyl handling rather than coordinated CoA-linked catabolism. Alongside these transformation-associated enzymes, methyl-accepting chemotaxis proteins and iron-sulfur cluster proteins comprised much of the associated signaling and electron-transfer functional complement.

### Ortholog-level functional mapping of gut microbial aromatic processing and redox defense

Proteome-wide KEGG Ortholog (KO) mapping further localized the aromatic-compound metabolic shift to the large intestine. To extend the curated protein analysis across the full metaproteome, all identified proteins were assigned to KOs and KEGG-anchored custom functional categories while retaining their genus-level attribution, enabling direct comparison of the functional architecture of each genus-resolved metaproteome between control and exposed conditions (**Fig. 3a–d; Supplementary Fig. 3a–f and 4a–r**). Notably, xenobiotic and aromatic compound metabolism showed the largest positive functional shift in the large intestine (weighted log2FC = +1.45), whereas such increase was not replicated in the small intestine (−0.31) (**Supplementary Fig. 3c,d**). Relatedly, condition-exclusive taxon-function profiles mapped the exposed-unique xenobiotic metabolic functions to uncultured *Acetatifactor*, *Lacrimispora*, *Eisenbergiella*, *Turicibacter*, *Roseburia* and *Kineothrix* in the large intestine (**Supplementary Fig. 3b,f**).

**Figure 3.**
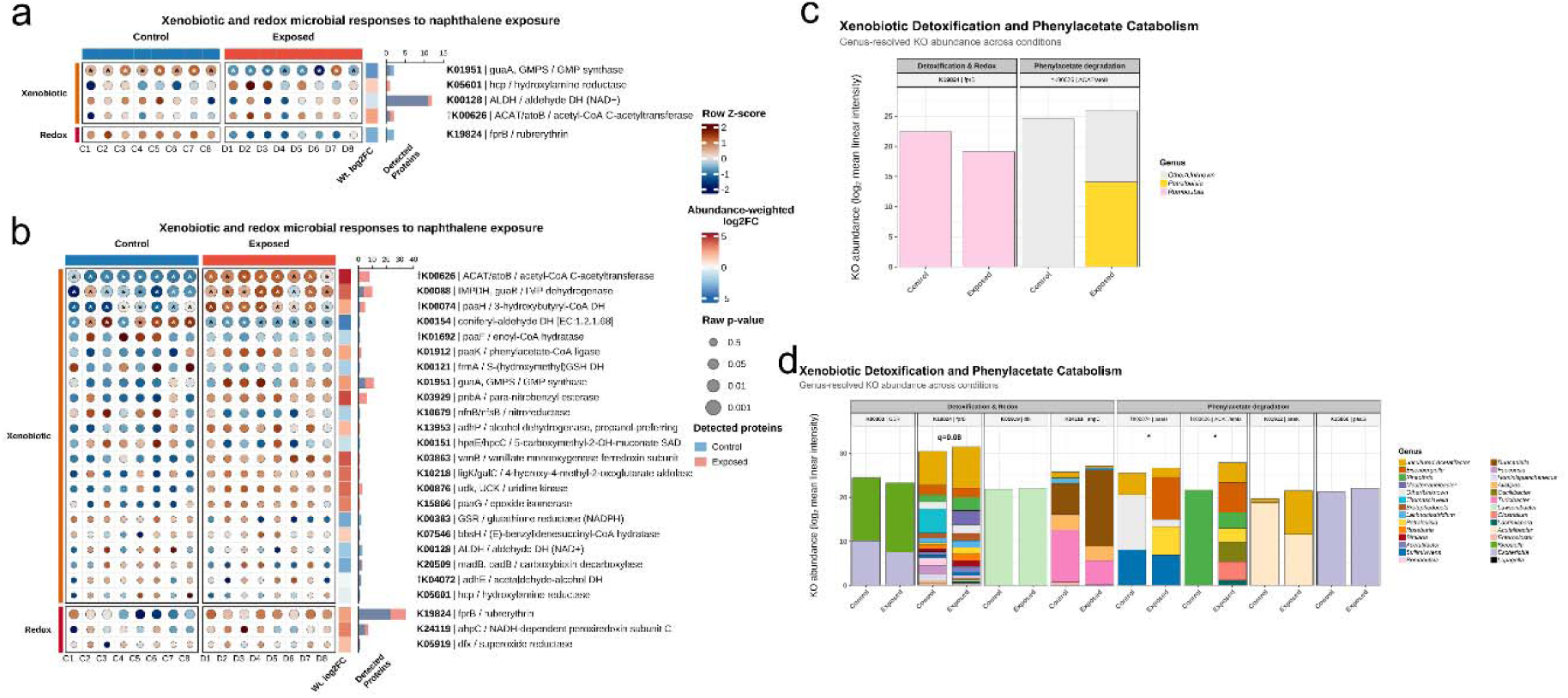
KO-level representation of aromatic/xenobiotic metabolism and redox defense in the intestinal metaproteome. (**a,b**) Profiles of aromatic/xenobiotic metabolic KOs in the small intestine (**a**) and large intestine (**b**) of control and naphthalene-exposed mice. Columns represent biological replicates (n = 8 per condition). Circle color denotes the row-wise Z-score of normalized KO abundance, and circle area encodes the raw Wilcoxon rank-sum P value. Asterisks indicate Benjamini–Hochberg-adjusted q < 0.05; daggers denote KO assignment to at least two curated functional classes. (**c, d**) Faceted stacked-bar aggregation of xenobiotic metabolic and redox functional shifts following PAH exposure. Each facet aggregates all KO-associated protein abundances belonging to the specified function into a single module-level abundance profile for exposed and control conditions of the small (**c**) and large (**d**) intestines. Bar heights depict module-level abundance computed as log_2_-transformed mean linear intensity across biological replicates (n=8): log_2_(mean linear intensity + 1). Bars are stacked by proportional taxonomic contributions to the total module abundance. Significance markers above facets indicate Wilcoxon rank-sum comparisons of total module abundance between control and exposed conditions after Benjamini-Hochberg correction (q < 0.05; trending, q < 0.10). “Other/Unknown” classification include proteins lacking confident genus assignment.

The distal KO profile depicted a broad aromatic-processing repertoire centered on phenylacetate-derived CoA intermediates (**Fig. 3b**). The abundance of 3-hydroxybutyryl-CoA dehydrogenase *paaH* (K00074) and acetyl-CoA C-acetyltransferase *ACAT/atoB* (K00626) increased significantly, while phenylacetate-CoA ligase *paaK* (K01912) and epoxide isomerase *paaG* (K15866) showed nominal increases. In addition, an increased vanillate monooxygenase ferredoxin subunit *vanB* (K03863) introduced an electron-transfer component of vanillate oxidation to protocatechuate, another aromatic-compound metabolic pathway. Moreso, exposure-increase of *ligK/galC* (K10218) linked the distal profile to terminal funneling of protocatechuate cleavage products into central carbon metabolism [50, 51], while paranitrobenzyl esterase *pnbA* (K03929) added a xenobiotic ester-hydrolysis function [52]. The large-intestinal metaproteome therefore displayed connected functions for aromatic activation, CoA-dependent intermediate processing and ring-product assimilation, along with ancillary xenobiotic transformation. In contrast, the proximal KO profile remained restricted and lacked an equivalent aromatic-processing architecture (**Fig. 3a,c,e**). Namely, hydroxylamine reductase *hcp* (K05601), representing nitrogen detoxification function [53] was detected only under exposure (q = 0.09) while *ACAT/atoB* (K00626) showed only a modest exposure-associated increase.

Remarkably, the distal increase in CoA-processing orthologs was accompanied by a broader redistribution of their parent organism. *Kineothrix* accounted for most K00626 abundance in controls, whereas under exposure its abundance was distributed across *Kineothrix*, *Lacrimispora*, *Oscillibacter*, *Clostridium*, *Petralouisia*, *Eisenbergiella* and uncultured *Acetatifactor* (**Fig. 3d**). K00074 showed a similar expansion, with prominent exposed contributions from *Petralouisia* and *Eisenbergiella*. Relatedly, ROS-defense functions showed taxonomic reorganization following naphthalene exposure (**Fig. 3f**). This was most observed in rubrerythrin (K19824) turnover, with the dominant control contribution from *Thomasclavelia* replaced under exposure by *Mediterraneibacter* and several other genera.

### Exposure-expansion of a hydrocarbon catabolic machinery in the gut

To resolve hydrocarbon-associated protein potential beyond the broad functional categories obtained through orthology-based annotation, detected metaproteins were queried against the CANT-HYD (Calgary approach to ANnoTating HYDrocarbon degradation genes) and HADEG (Hydrocarbon Aerobic DEgradation Genes) reference databases [54, 55] and classified by hydrocarbon substrate and pathway association (**Fig. 4a–f; Supplementary Fig. 5a,b**). Notably, hydrocarbon-catabolic protein annotations were detected in control and exposed metaproteomes from both intestinal compartments, with substantial increase following naphthalene exposure, particularly in the large intestine. Due to variation in the substrate specificity of individual hydrocarbon-marker assignments, subsequent mechanistic interpretation relied on convergent support across multiple related proteins rather than on any single annotation.

**Figure 4.**
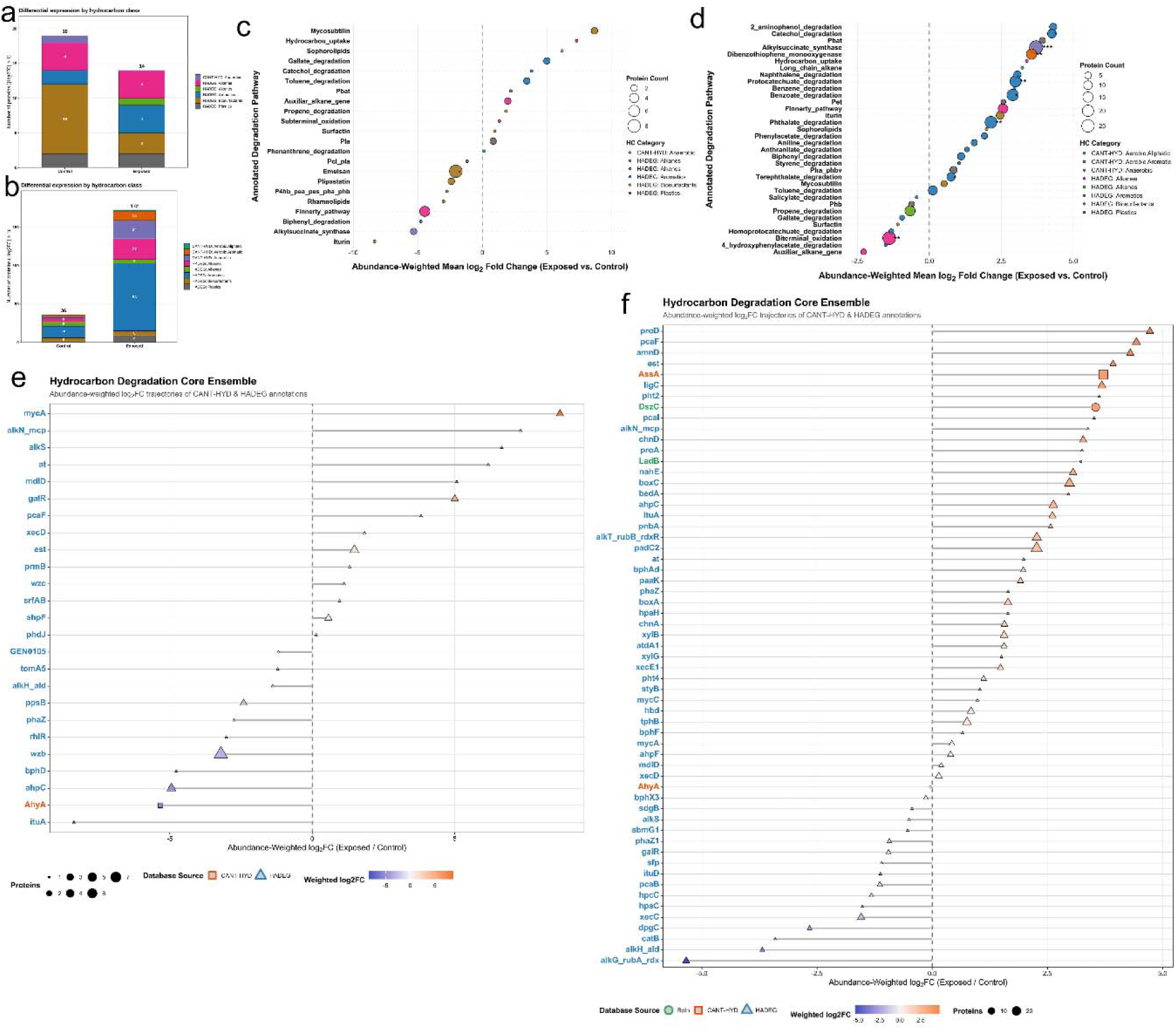
Hydrocarbon-focused annotation resolves compartment-specific catabolic potential in the intestinal metaproteome. (**a,b**) Distribution of hydrocarbon-associated protein annotations across control and naphthalene-exposed samples from the small intestine (**a**) and large intestine (**b**). Stacked bars depict the numbers of proteins assigned to hydrocarbon classes represented in CANT-HYD or HADEG. Values above bars indicate the total number of annotated proteins in each condition. (**c,d**) Abundance-weighted pathway-level profiles for CANT-HYD and HADEG annotations in the small intestine (**c**) and large intestine (**d**). Points show the abundance-weighted mean log2 fold-change for exposed relative to control samples. Positive values indicate greater representation under exposure and negative values indicate greater representation in controls. Point area denotes the number of proteins supporting each pathway annotation, and colour indicates the corresponding CANT-HYD or HADEG hydrocarbon class. Asterisks denote pathway-level one-sample Wilcoxon signed-rank tests of contributing protein log2 fold-changes after Benjamini–Hochberg correction, with q < 0.05 (*) and q < 0.01 (**). (**e,f**) Abundance-weighted fold-change profiles of individual hydrocarbon-degradation markers in the small intestine (**e**) and large intestine (**f**). Symbol shape identifies whether the annotation was recovered from CANT-HYD, HADEG or both databases; symbol area indicates the number of supporting proteins, and fill represents the abundance-weighted log2 fold-change (n = 8 replicates per condition).

Among proteins meeting the plotted ∣log_2_FC∣≥1 criterion, the number of hydrocarbon-associated annotations in the small intestine decreased from 19 in controls to 14 following exposure (**Fig. 4a; Supplementary Fig. 5a**). Though PAH relevant annotated hits decreased, the aromatic hydrocarbon assignments doubled, and an alkene-associated annotation was detected exclusively in the exposed condition. The net decrease was driven primarily by the reduction in biosurfactant-associated proteins from 10 to 3. Hence, the proximal response reflected redistribution toward aromatic-associated markers, which are chemically relevant to the administered naphthalene. In contrast, distal hydrocarbon-marker annotations increased 4.8-fold, from 36 in controls to 172 following exposure (**Fig. 4b**). Notably, the aromatic assignments rose from 15 to 88, a 5.9-fold increase, accompanied by 24 anaerobic and 12 aerobic-aromatic CANT-HYD assignments. Thus, the distal signature was chemically structured and ominated by proteins relating to aromatic hydrocarbon metabolism, alongside an anaerobic metabolic component that is consistent with its oxygen-restricted environment and chemistry.

Pathway-level analysis further separated the two compartmental hydrocarbon processing signatures (**Fig. 4c,d**). The small intestine retained a sparse and bidirectional profile. Marker sets associated with catechol, toluene and gallate degradation increased, whereas several other hydrocarbon-associated categories decreased (**Fig. 4c**). In contrast, the large intestine showed a broader positive profile that included marker sets assigned to naphthalene, catechol, protocatechuate, benzoate, benzene, biphenyl, dibenzothiophene and 2-aminophenol degradation (**Fig. 4d**).

Narrowing from pathway to protein-level trajectories identified the marker families underlying the aromatic-processing signatures (**Fig. 4e,f**). In the large intestine, positive shifts in *pcaI* and *pcaF* implicated CoA-transfer and thiolytic steps associated with downstream β-ketoadipate metabolism, while *paaK* marked entry into CoA-dependent phenylacetate processing [56–58]. Increased detection of *xylB* and *xylG* extended this repertoire to TOL-associated aromatic metabolism. Notably, *xylB* encodes benzyl alcohol dehydrogenase involved in upper-pathway aromatic oxidation, whereas *xylG* encodes 2-hydroxymuconic semialdehyde dehydrogenase, which processes an intermediate generated downstream of catechol meta-cleavage [59–62]. Hits assigned to *boxA* and *boxC* further implicated benzoyl-CoA-associated chemistry [63], while a *nahE*-model hit connected the protein ensemble to the naphthalene-associated pathway annotation. The recovery of *paaK* in both the KO-based and hydrocarbon-focused analyses provided an independent cross-annotation anchor (**Fig. 3b**; **Fig. 4f**). More so, the concordant *pca*, *paa*, *box* and *xyl* trajectories support increased representation of biochemically connected routes for aromatic-substrate processing in the exposed distal metaproteome.

### Distribution of the aromatic-catabolic machinery across gut commensals

Taxonomic attribution localized the exposure-associated hydrocarbon processing to distinct groups of gut commensal bacteria and showed that functional contribution did not scale directly with either total proteome representation or taxonomic abundance (**Fig. 5a–d; Supplementary Figs. 3a,b and 1e,f**).

**Figure 5.**
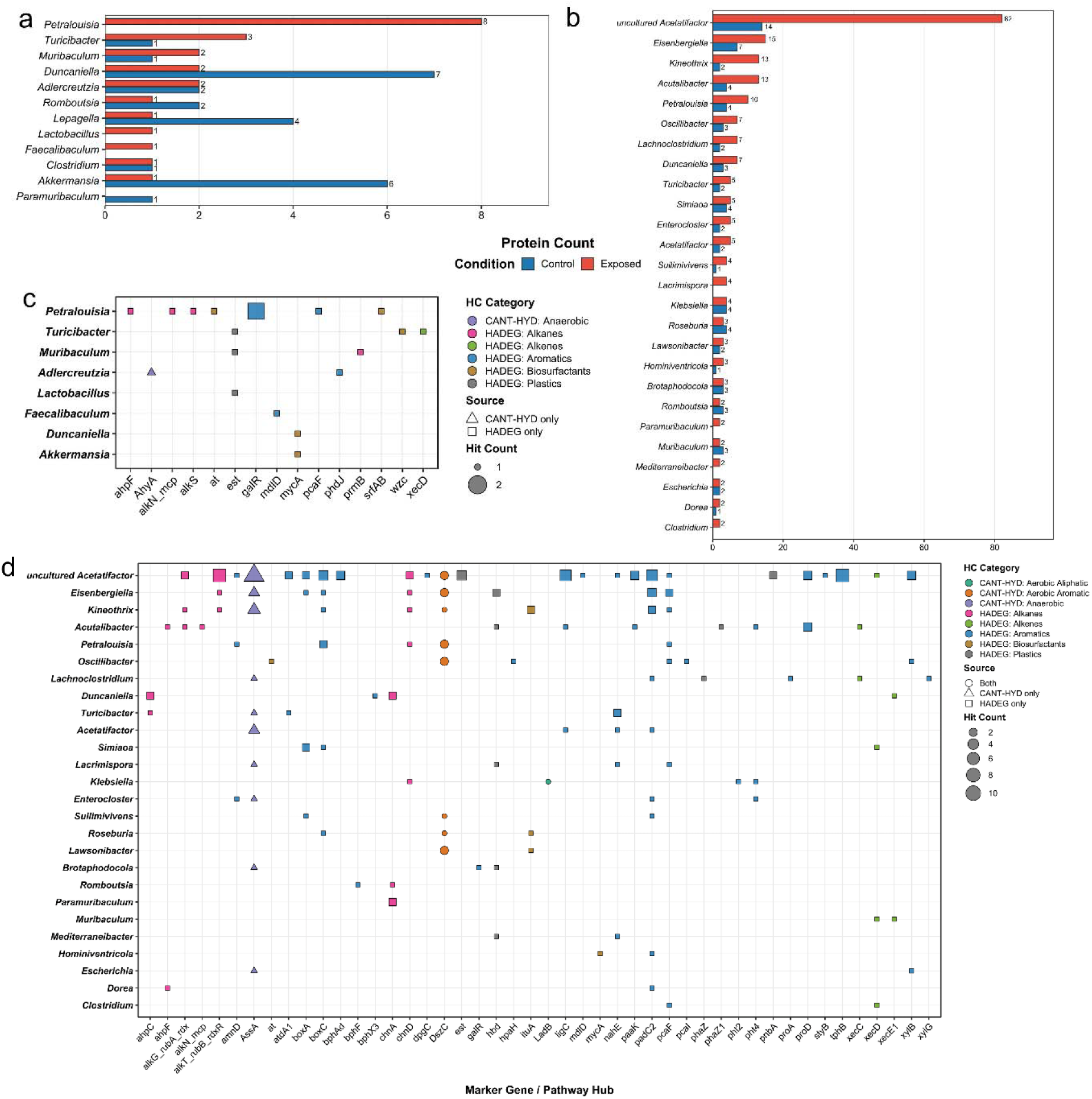
Genus-resolved attribution of putative hydrocarbon-catabolic proteins in the intestinal metaproteome. (**a,b**) Counts of hydrocarbon-annotated proteins among the top contributing microbial genera in the small intestine (**a**) and large intestine (**b**). Blue and red bars indicate proteins detected in control and exposed samples, respectively. Differences in the genus-level distribution of annotated proteins between conditions were evaluated using Monte Carlo χ² tests with 2,000 simulations. (**c,d**) Genus-by-marker hydrocarbon-catabolic attribution matrices for the small (**c**) and large (**d**) intestines. Rows denote genera and columns denote hydrocarbon-associated catabolic markers represented in CANT-HYD or HADEG. Point area indicates the number of distinct proteins assigned to each genus-marker combination. Point colour denotes the hydrocarbon annotation class, and point shape indicates whether the assignment was recovered from CANT-HYD, HADEG or both databases (n = 8 replicates per condition). Taxonomic assignments correspond to the genus annotations of the matched metaproteins.

In the small intestine, *Petralouisia* depicted eight exposed-associated marker proteins, emerging as the leading proximal contributor to hydrocarbon metabolism, with additional contributions from *Turicibacter* and *Muribaculum* (**Fig. 5a**). On the other hand, the large intestine showed a broader exposure-associated mobilization, with the largest increase in an uncultured *Acetatifactor*, which increased from 14 to 82 proteins (**Fig. 5b**). This was accompanied by *Eisenbergiella*, which increased from 7 to 15 proteins, while *Kineothrix* and *Acutalibacter* each increased to 13 and *Petralouisia* increased from 4 to 10. Additional exposure-associated increases were distributed across *Oscillibacter, Lachnoclostridium, Duncaniella, Turicibacter, Enterocloster* and *Acetatifactor*. Moreover, *Lacrimispora*, *Paramuribaculum*, *Mediterraneibacter* and *Clostridium* contributed hydrocarbon-associated proteins only under exposure, depicting a largely inductive response. The taxonomic distribution differed strongly between conditions (Monte Carlo χ², B = 2,000, p < 0.001), consistent with amplification of a broadly baseline-detected distal protein set together with an exposure-added component. Strikingly, hydrocarbon-associated representation across different genera was not proportional to the magnitude of their expressed proteome. Notably, *Kineothrix* and *Acutalibacter* had more hydrocarbon-associated annotations than *Petralouisia* despite their relatively smaller detected proteomes. Similarly, *Lactobacillus* which had one of the top-most exposure-associated proteomic representations across both intestinal segments, depicted minimal hydrocarbon-associated protein representation (**Fig. 5b; Supplementary Fig. 3a,b and 5a,b**). These contrasts indicate that naphthalene exposure mobilized hydrocarbon-associated proteins within a functionally specialized subset of gut commensals.

Marker-level attribution further resolved the taxonomic structure of the hydrocarbon-associated proteome (**Fig. 5c,d**). In the small intestine, *Petralouisia*, depicted a wide array of exposure-associated markers including aromatic markers *pcaF* and *galR* (**Fig. 5c**). The large intestine showed a broader and more chemically diverse attribution pattern (**Fig. 5d**). An uncultured *Acetatifactor* depicted the most annotation profile, with assignments across alkane-associated markers, anaerobic hydrocarbon markers, naphthalene pathway marker *nahE*, and several other aromatic-markers: *paaK*, *pcaF*, and *xylB*. The *nahE* marker, which encodes a hydratase-aldolase that converts a ring-opened naphthalene intermediate into salicylaldehyde and pyruvate [64], was also encoded by *Turicibacter*, *Acetatifactor*, *Lacrimispora*, and *Mediterraneibacter*.

### Chemoproteomics characterization of naphthalene-engaged gut microbial proteome

To move beyond protein expression and directly capture proteins that physically engage PAH-like chemical structures, we employed a targeted chemoproteomics workflow. We synthesized two resin-immobilized naphthalene probes, **Br-NAP** and **DADPS-BrNAP**, each containing a naphthalene recognition element and a chloroacetamide capture moiety, together with an acetyl-capped control resin (**CR-C**) (**Fig. 6a–c**; **Scheme S1 and S2**). Probe performance was first evaluated in the canonical naphthalene degrader *Pseudomonas putida G7*. Relative to **CR-C**, both probes enriched several naphthalene-associated proteins (**Supplementary Fig. S6a–h**). However, **Br-NAP** showed the strongest selectivity for aromatic hydrocarbon-associated proteins, with aromatic-class proteins forming the largest category of significantly enriched targets and exceeding alkane, alkene and biosurfactant classes (**Fig. 6d; Supplementary Fig. S6f,h**). Several established naphthalene-catabolic markers were among the most enriched proteins, including *nahE*, *nahF* and the naphthalene dioxygenase regulatory marker *ndoR* [65–67]. These data established **Br-NAP** as the more selective probe for enriching proteins associated with defined naphthalene metabolism in a positive-control organism.

**Figure 6.**
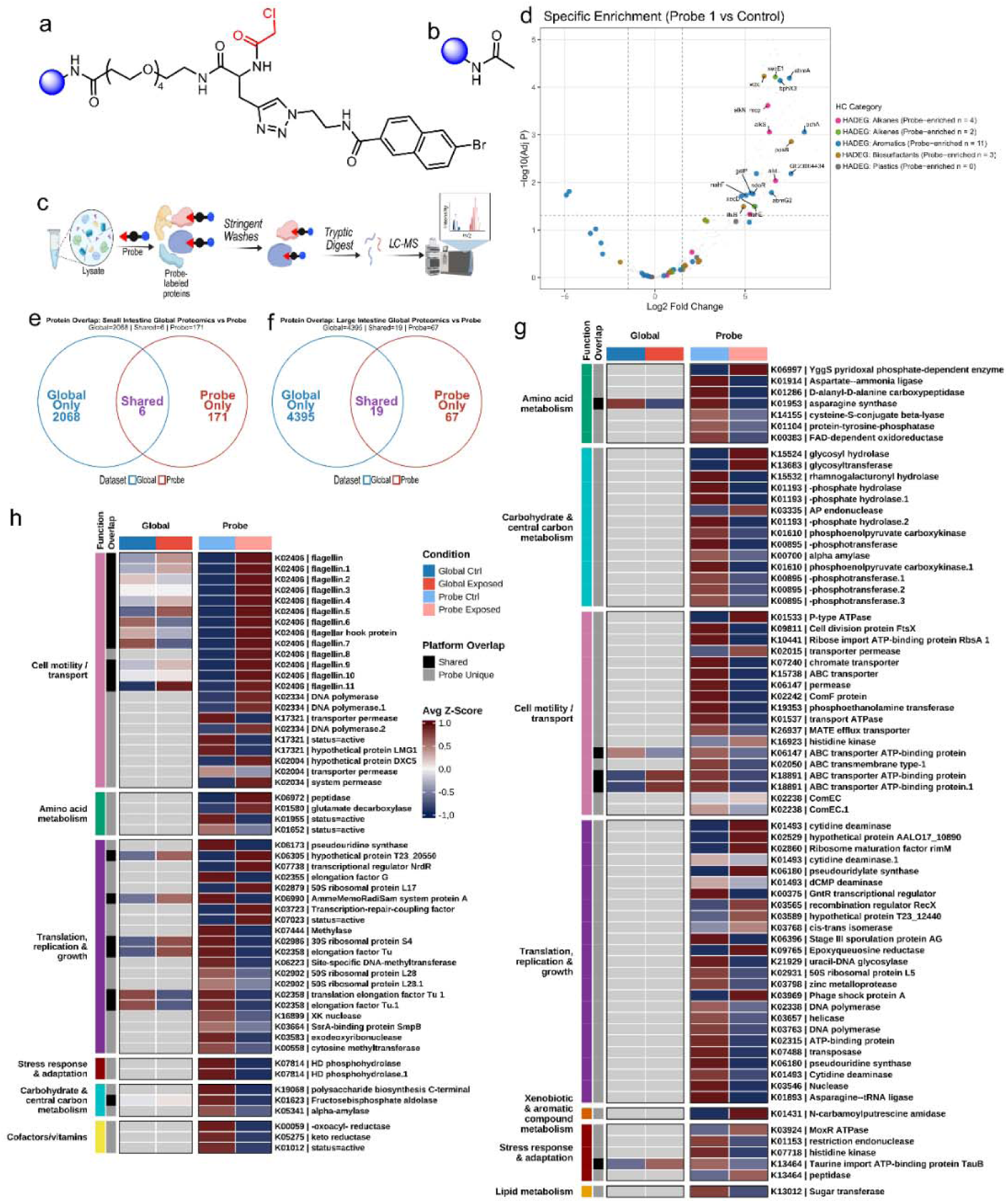
Chemoproteomics profiling uncovers the naphthalene-probe-engaged proteome. (**a,b**) Structures of the resin-immobilized naphthalene probe (**Br-NAP**) featuring a chloroacetamide (red) reactive moiety (**a**) and the non-reactive control probe, **CR-C** (**b**). (c) Schematic of the resin-dependent chemoproteomic workflow. (d) Volcano plot highlighting **Br-NAP** probe-enriched hydrocarbon catabolic proteins in *P. putida* G7 (**e,f**) Venn diagram comparing probe-enriched proteins from the chemoproteomic dataset with the proteins quantified in the global metaproteome of small (**e**) and large (**f**) murine intestinal samples. (**g,h**) Heatmap of all-probe-enriched proteins and globally abundant proteins in exposed and control sample groups of the small (**g**) and large (**h**) intestines. Columns are ordered by global control, global exposed, probe-labeled control, and probe-labeled exposed sample groups. Rows correspond to individual probe-enriched proteins, colors represent row-scaled Z-scores of mean protein intensities. Grey cells correspond to proteins that were below the limit of detection in the global metaproteomic analyses but were enriched by **Br-NAP** probe.

Importantly, application of **Br-NAP** to murine intestinal metaproteomes recovered a protein set that was largely distinct from the discovery metaproteome. In the small intestine, only six proteins were shared between the global and probe-enriched datasets, whereas 171 proteins were recovered only by the probe (**Fig. 6e**). In the large intestine, 19 proteins were shared and 67 were probe-exclusive (**Fig. 6f**). These observations are consistent with enrichment of low-abundance and chemically accessible proteins that were undetectable to global metaproteome profiling.

Functional annotation showed that the probe-engaged proteome was segment-specific (**Fig. 6g,h; Supplementary Fig. S7a,b**). In the small intestine, **Br-NAP** enriched transporters, central-metabolic proteins and nucleotide-associated enzymes, including cytidine deaminase (K01493), as well as β-ureidopropionase (K01431), and a transporter permease (K02015). Stress and defense-associated proteins were also represented, including an exposure-enriched MoxR-family AAA+ ATPase (K03924) and a baseline restriction endonuclease (K01153). On the other hand, the large-intestine probe-enriched proteome was dominated by motility-associated proteins. Flagellin (K02406) was the most frequently represented ortholog, with 13 probe-enriched proteins spanning a flagellin series and associated flagellar assembly component (**Fig. 6h; Fig. S9b**). Beyond motility, **Br-NAP** enriched glutamate decarboxylase (K01580), the committed enzyme of the glutamate-dependent acid-resistance system and a producer of gamma-aminobutyric acid, together with oligopeptide transport and translation-associated proteins.

## Discussion

PAH toxicology is generally interpreted through host absorption and biotransformation, although ingested compounds also encounter a metabolically active intestinal microbiome. By integrating 16S rRNA gene sequencing, global metaproteomics, chemoproteomics, and specialized hydrocarbon database annotation, our study identifies a hydrocarbon-associated protein architecture that was concentrated in the large intestine and poorly predicted by community composition. Notably, exposure explained 48.4% of distal metaproteomic variation but only 7.9% of the corresponding Bray-Curtis compositional variation, whereas the difference between these layers was narrower proximally. These observations also extend the taxon-function decoupling principle [68, 69] by showing that the principal microbial contributors to hydrocarbon-associated protein representation were largely invisible to 16S rRNA gene differential-abundance profiling. More so, this distal metabolic concentration is consistent with prior experimental evidence that colonic microbiota can transform naphthalene into biologically active products [20] though the responsible enzymes therein were not characterized. Our study extends the observation by resolving the enzyme-level context for this spatial pattern of hydrocarbon-processing in the gut (**Fig. 7**).

**Figure 7.**
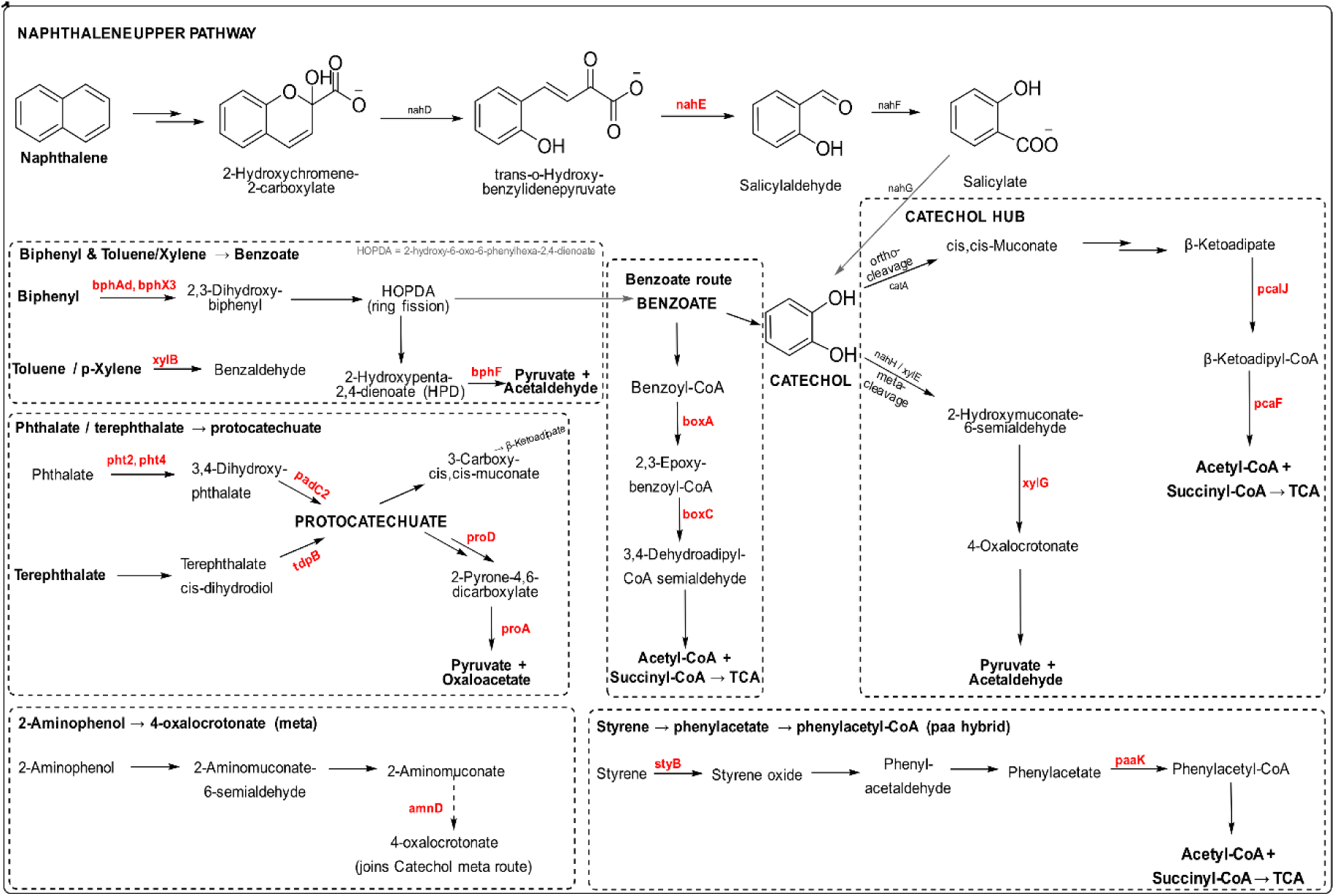
Hydrocarbon-catabolic signatures from gut commensals converge across aromatic-processing pathways. Schematic integration of CANT-HYD- and HADEG-annotated proteins detected in the gut metaproteome with established routes of aerobic and anaerobic aromatic-compound metabolism. Enzyme and marker-gene labels shown in red denote proteins detected in this study. The mapped proteins occupy several levels of aromatic processing, including upper-pathway transformations, ring-cleavage-associated reactions and downstream funneling through benzoate, catechol, protocatechuate, phenylacetyl-CoA and 4-oxalocrotonate-associated routes. Representative detected markers include the naphthalene-associated enzyme ***nahE***; biphenyl-associated ***bphAd***, ***bphX3*** and ***bphF***; benzoate-CoA pathway components ***boxA*** and ***boxC***; catechol- and protocatechuate-associated markers ***xylG***, ***pcaI*** and ***pcaF***; and the phenylacetate-activating enzyme ***paaK***.

A clear exposure-associated separation in biochemical function was observed between intestinal compartments. The small intestinal profile showed a sparse hydrocarbon-processing machinery largely represented by select catechol-, toluene- and gallate-associated marker sets. By contrast, the large-intestinal architecture converged on sequential reactions characteristic of activated aromatic-intermediate processing (**Fig. 7**). Acyl-CoA dehydrogenases, enoyl-CoA hydratases, hydroxyacyl-CoA dehydrogenases and thiolases increased alongside electron-transfer flavoproteins and redox-defense proteins, while the KO and hydrocarbon-focused analyses narrowed this broader pattern to *paa*, *pca, box* and *xyl* marker families. Recovery of these functions across the protein-level heatmap, KO mapping and hydrocarbon-focused annotation provides convergent support for CoA-dependent aromatic-intermediate processing as a central feature of the distal PAH-associated metaproteome. This architecture is biochemically consistent with the bacterial phenylacetate catabolic pathway, a hybrid route in which *paaK* activates phenylacetate to phenylacetyl-CoA, *paaABCDE* epoxidizes the aromatic ring, and *PaaG* and *paaZ* mediate rearrangement and hydrolytic ring opening [70, 71]. The resulting CoA-linked intermediates are subsequently processed through *paaF*-, *paaH*- and *paaJ*-mediated hydration, dehydrogenation and thiolysis, ultimately yielding acetyl-CoA and succinyl-CoA [71, 72]. Thus, the distal enrichment of hydratases, hydroxyacyl-CoA dehydrogenases, thiolases and related CoA-processing proteins mirrored these downstream reaction classes, while recovery of *paaK*, *paaG* and *paaH* provided pathway-specific anchors. Though not all canonical components were detected, the combined evidence supports a coherent partial architecture for phenylacetate-derived CoA processing.

Beyond phenylacetate metabolism, the combined *pca*, *xyl*, *box* and *nahE*-model assignments (**Fig. 7**) suggest that the distal metaproteome was organized around shared biochemical transitions that connect aromatic oxidation and ring processing to CoA-linked downstream metabolism. This organization is consistent with the funneling logic of bacterial aromatic catabolism, in which chemically distinct substrates converge on a restricted set of ring-cleavage products and activated intermediates before entering central carbon metabolism [73]. The *nahE*-model hit extends this architecture to an upper-pathway reaction associated with naphthalene [64], whereas *pca*, *xyl* and *box* functions support complementary downstream processing steps. Together, these support a distributed distal repertoire that links naphthalene-associated and related aromatic transformations to convergent CoA-dependent processing.

This convergent architecture also included proteins assigned to anaerobic hydrocarbon metabolism, placing the distal signal within a biochemical context appropriate to the oxygen-limited colon. In characterized anaerobic naphthalene pathways, substrate activation can proceed through carboxylation to 2-naphthoate, CoA ligation and reductive dearomatization of 2-naphthoyl-CoA [27, 74]. Although the present metaproteome does not establish this route or a complete 2-naphthoyl-CoA pathway, the combined detection of anaerobic CANT-HYD markers, CoA-processing proteins and low-potential electron-transfer functions is consistent with such anaerobic aromatic catabolic paradigms. The concomitant detection of aerobic markers also aligns with the radial oxygen gradient of the colon, where relatively oxygenated mucosal regions coexist with an anaerobic lumen [31, 75]. This spatial heterogeneity could support distinct aromatic-processing chemistries across the colon, with oxygen-dependent and anaerobic functions partitioned among different microenvironments or microbial taxa.

Taxonomic attribution localized the highest hydrocarbon-associated protein complements to a restricted set of distal gut families. Four of the prominent genera, an uncultured *Acetatifactor*, *Kineothrix*, *Petralouisia*, and *Eisenbergiella* belong to Lachnospiraceae, while *Acutalibacter* belongs to Oscillospiraceae [76–78]. This concentration is notable because these families are prevalent in the human gut [79] and are best known for anaerobic carbohydrate fermentation and short-chain fatty acid production [80, 81]. The detection of hydrocarbon-associated proteins in these taxa therefore extends such annotation to native gut commensals not otherwise linked to this chemistry. An uncultured *Acetatifactor* was the dominant distal contributor, whereas *Kineothrix* and *Acutalibacter* allocated a larger fraction of their substantially smaller proteomes to hydrocarbon-associated functions, compared with *Lactobacillus*, *Lacrimispora* and *Mediterraneibacter*. The latter taxa underwent broad exposure-associated proteomic expansion but contributed comparatively few hydrocarbon markers, suggesting that their response centered more on stress tolerance, substrate acquisition and fermentative remodeling. *Acutalibacter*, by contrast, showed substantial exposure-associated hydrocarbon representation despite a control-directed 16S rRNA gene profile. Thus, the aromatic-processing repertoire emerged as a genus-restricted feature embedded within clades that are also prominent in humans and did not directly scale with taxonomic abundance or total proteome depth. This organization explains why taxonomic abundance failed to predict the response and suggests that interindividual differences in the carriage and expression of these genera could modify intestinal PAH fate in humans.

The hydrocarbon-specific analysis addressed a major challenge in determining the bacterial proteins interacting with and metabolizing PAHs, overcoming limitations of standard functional annotation. Applying the specialized HADEG and CANT-HYD hydrocarbon databases resolved this blind spot, revealing an extensive, exposure-associated hydrocarbon-degradation proteome in the gut (**Fig. 7**). HADEG contributed marker families from experimentally validated aerobic degradation sequences [54], and CANT-HYD added phylogeny-derived models that include anaerobic hydrocarbon-activation pathways [55]; hence, their complementary use recovered broader aromatic and anaerobic hydrocarbon degradation repertoires. While these markers indicate sequence-based potential rather than confirmed in vivo catalysis, applying this paired database strategy successfully unmasked both latent and exposure-induced, pathway-resolved aromatic repertoire that broad annotations systematically misassign.

Chemoproteomics further resolved a distinct, accessibility-defined layer of the naphthalene-associated proteome. In *P. putida* G7, the **Br-NAP** probe enriched established naphthalene-pathway proteins, including *nahE*, *nahF* and *ndoR* [65–67], validating the probe scaffold in a canonical degrader. In the gut, however, the aromatic-catabolic proteins identified by global metaproteomics and hydrocarbon-focused annotation did not meet the enrichment threshold. Nonetheless, the probe-enriched proteins were largely absent from the metaproteomics dataset and were dominated by transport, stress-response and surface-associated proteins, including distal flagellins. This divergence distinguishes protein representation from chemical accessibility. More so, **Br-NAP** probe reports engagement with a resin-immobilized naphthalene scaffold, not substrate turnover. Failure to enrich the annotated catabolic proteins may reflect their limited active-site accessibility or altered ligand presentation after immobilization. However, the flagellin and glutamate decarboxylase enrichments nominate possible gut-PAH exposure links to TLR5 sensing and acid-resistance or GABA-associated physiology [82–85]. Overall, the largely distinct chemoproteomic protein enrichment shows that probe accessibility constitutes an orthogonal property of the gut proteome that cannot be inferred from discovery-level abundance alone.

## Conclusion

Native gut commensals express a previously unresolved aromatic-hydrocarbon catabolic proteome, and naphthalene exposure mobilized this machinery most strongly in the large intestine. Independent annotation frameworks converged on upper naphthalene-associated markers and proteins linked to phenylacetate, β-ketoadipate, catechol and benzoyl-CoA processing, alongside anaerobic hydrocarbon markers and downstream CoA-linked chemistry. These proteins were distributed across human prevalent Lachnospiraceae and Oscillospiraceae families, including an uncultured *Acetatifactor*, *Kineothrix*, *Acutalibacter*, *Eisenbergiella*, *Petralouisia* and other native commensals, indicating a community-distributed catabolic machinery. Most of these taxa are known primarily for fermentation, carbohydrate utilization or host metabolic effects, with no currently established role in hydrocarbon catabolism. The recovery of hydrocarbon-associated proteins from these commensals therefore extends aromatic-processing potential into gut lineages not conventionally linked to this chemistry. Notably, this exposure-associated catabolic representation was not predicted by 16S rRNA gene abundance or total proteome size, showing that the intestinal response to naphthalene was primarily functional and spatially organized. Chemoproteomics identified a largely separate naphthalene-engaged proteome, further distinguishing predicted function and protein abundance from chemical engagement. Critically, these provide an enzyme-level map of latent and exposure-amplified hydrocarbon-processing potential in native gut commensals. Together, these findings provide a functional framework for intestinal PAH metabolism and identify the distal microbiome as a previously underrecognized reservoir of aromatic-processing potential. Determining whether this repertoire promotes naphthalene detoxification, bioactivation or both will require isotope-resolved substrate tracing, metabolite toxicokinetics, biochemical characterization of prioritized proteins and probes designed for soluble or mechanism-based engagement.

## Supporting information

Metaproteomics data

Large Intestine Functional Analysis

Small Intestine Functional Analysis

16s rRNA squencing data

Chemoproteomics data

## Supporting Information

The Supporting Information is available free of charge on the Publications website.

Supplementary text and supplementary figures S1 through S7 with accompanying legends, Schemes S1 and S2, and chemistry synthetic methods and characterization. (PDF)

Supplementary File 1: S1 -Metaproteomics crosstab, differential abundance metrics and protein-level statistical results and figure source data. (XLSX)

Supplementary File 2: S2 -Large intestine functional analyses figure source data. KO, CANT-HYD and HADEG reference tables are appended. (XLSX)

Supplementary File 3: S3 -Small intestine functional analyses figure source data. KO, CANT-HYD and HADEG reference tables are appended. (XLSX)

Supplementary File 4: S4 -Processed 16S rRNA data, figure source data, and statistical output files. (XLSX)

Supplementary File 5: S5 -Chemoproteomics crosstab, differential abundance metrics and protein-level statistical results and figure source data. (XLSX)

## Ethics approval and consent to participate

All animal experiments and experimental protocols were reviewed and approved by the Institutional Animal Care and Use Committee (IACUC) of Baylor University under protocol number [#2355412-2]. All mouse handling, sub-acute oral gavage exposures, and tissue collection procedures were performed in strict compliance with the National Institutes of Health Guide for the Care and Use of Laboratory Animals. This study did not involve human participants, human data, or human tissue samples.

## Availability of data and materials

The datasets generated and analyzed during the current study are available in public, community-recognized repositories. The 16S rRNA gene amplicon sequencing raw fastq reads have been deposited in the National Center for Biotechnology Information (NCBI) Sequence Read Archive (SRA) and are accessible under the BioProject accession number [PRJNA1484644].

The raw mass spectrometry files, database search parameters, and label-free quantitative output tables for both the global discovery metaproteomics and the activity-based chemoproteomics datasets have been deposited to the MassIVE repository under the dataset identifier MSV000102316, password i2PCY922s40g3CH1.

The custom R processing pipelines, statistical scripts utilizing the limma framework, ontology assignment rules for environmental catabolic markers, and figure generation loops are fully documented and publicly accessible via our laboratory GitHub repository at https://github.com/WrightLab-Baylor.

## Acknowledgments

This work was supported by the James R. Schofield Endowment in Biomedical Science to A.T.W. The authors thank the staff of the Baylor University Vivarium facility for their assistance with mice husbandry. We also acknowledge the core staff at the Baylor University Mass Spectrometry Center, Nuclear Magnetic Resonance Facility, and Molecular Biosciences Center for their technical support. We thank the Gunsch Laboratory at Duke University for kindly providing the *P. putida* G7 strain. We are also grateful to Abhijit Maji (Ph.D) for providing insightful suggestions and editorial assistance during the preparation of this manuscript. Table of Contents Graphic was generated with Biorender. Nwike, C. (2026) https://BioRender.com/r82v505.

## For Table of Contents Only

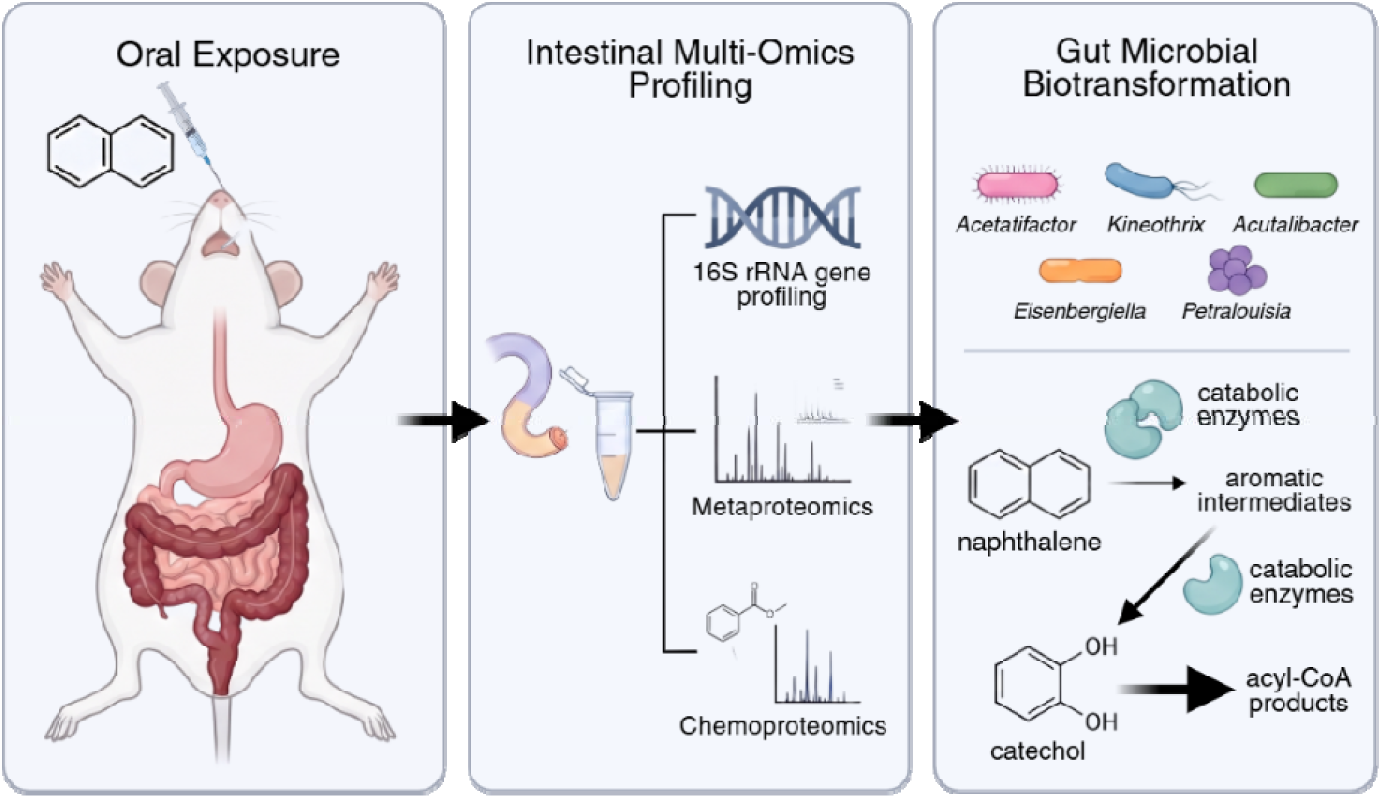

