## Supplemental Information for "Commensal Bacteria in the Gut Microbiome Encode Previously Unrecognized Aromatic Hydrocarbon Catabolic Machinery"

Figure S1…………………………………………………………………………………………………….9

Figure S2 …………………………………………………………………………………………………..10

Figure S3 …………………………………………………………………………………………………..11

Figure S4 …………………………………………………………………………………………………..12

Figure S5 …………………………………………………………………………………………………..13

Figure S6 …………………………………………………………………………………………………..14

Figure S7 …………………………………………………………………………………………………..15

S4. Synthetic Details ……………………………………………………………………………………………….16

S5. Synthetic Schemes to Access Resin-immobilized Naphthalene Probes ………………………………..18

S6. NMR Spectra …………………………………………………………………………………………………...25

Supplementary References ……………………………………………………………………………………….27

**Supplementary Methods**

**S1. Murine gut contents and *Pseudomonas putida* G7 cells preparation for global proteomics.**

***Pseudomonas putida G7* cultivation, naphthalene catabolic induction, and protein extraction.** *P. putida* G7 was utilized as a model naphthalene-degrading bacterium to serve as an induced functional reference for global and chemoproteomic hydrocarbon degradative pathway mapping. Primary seed cultures were grown by inoculating 2.0 mL of Glucose-Tryptone-Yeast extract (GTY) medium with *P. putida* G7 cells, followed by overnight incubation at 28 ºC under constant agitation. The overnight culture was then expanded via a 1:75 dilution into 150 mL of modified medium where naphthalene served as the primary inducing carbon source (Naphthalene-Tryptone-Yeast extract medium). Subsequently, the culture was grown at 28 ºC with constant agitation until an optimal mid-logarithmic optical density (OD_600_) of 0.7. Bacterial cells were harvested by centrifugation at 3,600 x *g* at 23 ºC for 20 min. The resulting cell pellets were washed twice with 20 mL of 1x phosphate-buffered saline (PBS) buffer, pelleting each time by centrifugation at 3,600 x g at 23 ºC for 5 min. Cell pellets were then resuspended in 1 mL of ice-cold 1x phosphate-buffered saline (PBS) and aliquoted into 2 mL screw top microtubes pre-loaded with 100 μL of 0.1 mm silica zirconium beads for cell lysis. Lysing was carried out on Omni Bead Ruptor Elite with bead beater settings: 4.20 m/s, 45 s, 3 cycles. Samples were transferred to ice for 5 min between cycles. Post-lysing, the microtubes were centrifuged at 17,000 x *g*, 4 ºC, 30 min. Supernatant were aliquoted into 1.5 mL safe-lock tubes, and standard bicinchoninic acid (BCA) assay was performed to determine total protein concentrations.

**Metaproteomics sample isolation and protein extraction.** Murine gut contents, isolated from the small and large intestine segments were suspended in ice-cold PBS (2.0 mL) and subjected to intermittent 30 s vortexing and ice-incubation to achieve complete rehydration and homogeneity. Fully homogenized samples were then centrifuged at 1,000 x *g*, 4 ºC, 10 min and supernatant were carefully collected into 2 mL screw top microtubes pre-loaded with 200 μL of 0.1 mm silica zirconium beads for cell lysis. Lysing was carried out on Omni Bead Ruptor Elite with bead beater settings: 4.20 m/s, 45 s, 3 cycles. Samples were transferred to ice for 5 min between cycles. Post-lysing, the microtubes were centrifuged at 17,000 x *g*, 4 ºC, 30 min and supernatant carefully isolated. For protein concentration quantification, dual 50 μL aliquots were drawn from each supernatant and transferred to independent 1.5 mL microcentrifuge tubes. Protein precipitation was initiated by adding 200 μL of ice-cold acetone to each tube. The solutions were briefly vortexed and incubated at -20 ºC for 30 min, before precipitated protein pellets were harvested by centrifugation at 18,213 x *g*, 4 ºC, 10 min The supernatant was decanted and protein pellets dried at room temperature for 30 min for complete acetone evaporation. Subsequently, dried pellets were resuspended in Milli-Q H_2_O and BCA assay performed to determine total protein concentrations.

**Protein reduction, alkylation, and tryptic digestion.** Soluble protein fractions from both the large and small murine intestinal contents (n = 8 replicates per exposed or control group) and *P. putida* G7 (n = 3 replicates) were normalized to 100 µg of total protein in 50 mM NH_4_HCO_3_, pH 8.0. Proteins were denatured by the treatment of samples with solid urea (75 mg) to give a final concentration of ~8M urea. Following this, reduction of protein disulfide bonds was carried out by the addition of 3 µL of freshly prepared 1 M dithiothreitol (DTT) in Milli-Q H_2_O to samples, with 45 min incubation on a thermomixer at 50 ºC and 1200 rpm. Reduced samples were cooled to room temperature for 5 min. Subsequently, alkylation was performed by the addition of 10 µL of 200 mM iodoacetamide (IAA) prepared in 50 mM NH_4_HCO_3_ (pH 8.0), followed by sample incubation and mixing under foil (rt, 45 min). Additional DTT (3 µL of 500 mM) was added to samples along with incubation at 30 ºC for 30 min with agitation at 1200 rpm to scavenge excess IAA. Next, samples were diluted with the addition of 320 µL of 50 mM NH_4_HCO_3_ (pH 8.0) to reduce urea concentration and 1 µL of 1 M CaCl_2_ were also added to prime samples for optimal trypsinization. Protein digestion was then performed with the addition of sequencing-grade porcine trypsin, pre-activated by incubation at 37 ºC for 10 min in manufacturer-supplied buffer. Trypsin was added at a ratio of 1 µg trypsin per 50 µg total protein to all samples and incubated at 37 ºC while shaking overnight (~16 h) at 1200 rpm. Following this, peptide mixtures were desalted using reversed-phase C18 solid-phase extraction (SPE) cartridges. Eluted peptides were then concentrated to dryness in a speed vacuum and stored at -20 ºC.

Prior to LC-MS analysis, dried peptides were resuspended in 40 μL MS resuspension buffer (0.1% formic acid, 98% H_2_O, and 2% acetonitrile solution) and heated to 37 ºC for 5 min with 1000 rpm shaking. The solution was then transferred to ultracentrifuge tubes (Beckman-Coulter 5/16” × 1 3/8”) and centrifuged at 53000 x *g* for 20 min at 4 ºC. Supernatant was carefully drawn without disturbing the pellets and transferred to low-retention microcentrifuge tubes. Subsequently, sample peptide concentrations were determined using a fluorometric peptide quantification assay, according to the manufacturer’s instructions, and normalized to 0.25 µg/µL with the MS resuspension buffer. Normalized samples (30 µL) were individually transferred into 0.2 mL amber TPX screw-cap 9 mm vials with short-thread conical glass inserts, sealed and stored at -20 ºC till LC-MS/MS analyses.

**S2. Resin-based probe labeling and chemoproteomic analyses of microbial proteomes.**

**On-resin probe labeling and LC-MS/MS chemoproteomic profiling of *P. putida* G7 and gut microbial proteomes.** Lysates of either *P. putida* G7 (n = 4 replicates) or the large and small murine intestinal contents (n = 8 replicates per exposed / control group) were normalized to 500 μg total protein concentration in PBS. Probe labeling proceeded with the incubation of protein lysates with 5 μL DMSO slurry of either the 100 μM resin-ABP (**Br-NAP**, **DADPS-Br-NAP**) or the acetyl-capped resin control probe (**CR-C**) (**Fig. 6a–c**) for 90 min at 37 ºC with gentle agitation. Resin beads were then collected by centrifugation (1 min at 600 x *g*) and resuspended in 1 mL of 4% SDS in PBS for protein denaturation. Subsequently, samples were incubated with rotation for 1 h at 37 ºC. Beads were then pelleted and washed 3x sequentially with: 1.0 mL of 1X PBS, 1% SDS in PBS, 6M urea in 25 mM NH_4_HCO_3_ (pH 8), 1X PBS, Milli-Q H_2_O, and 25 mM NH_4_HCO_4_ (pH 8) to deplete unbound and non-specifically interacting proteins. For each wash, 1.0 mL of washing solution was added to beads, gently vortexed, and centrifuged (600 x *g*, 1 min) with careful removal of the supernatant. After the final wash, protein-bound beads were transferred to 1.5 mL microcentrifuge tubes.

**On-bead protein reduction, alkylation, and digestion.** Enriched proteins on the resin-bound probe were resuspended in 400 μL 6 M urea (freshly prepared in 25 mM NH_4_HCO_3_, pH 8.0). The proteins were reduced via the addition of 20 μL of 100 mM TCEP, with thermal incubation and agitation at 50 ºC for 50 min. Subsequently, alkylation was performed by the addition of 20 μL of 200 mM iodoacetamide to the samples, followed by incubation for 45 min at room temperature, under foil. Beads were pelleted and washed 8x with 1 mL 1X PBS, followed by four 1 mL 25 mM NH_4_HCO_3_ (pH 8) washes. Beads were collected by centrifugation, resuspended in 200 μL of 25 mM NH_4_HCO_3_ (pH 8) and 0.125 μg trypsin was added and incubated overnight at 37 ºC with 1200 rpm shaking. Samples were then centrifuged (10,500 x *g*, 5 min) and supernatant collected and transferred to new individually wrapped tubes. Beads were resuspended in additional 150 μL of 25 mM NH_4_HCO_3_, incubated at 37 ºC for 10 min with 1,200 rpm agitation. Following centrifugation, supernatant was collected and combined with prior collections. The tubes were covered with breathable membranes and transferred to a speed vacuum to evaporate liquids. Samples were resuspended in 40 μL MS resuspension buffer (0.1% formic acid, 98% H_2_O, and 2% acetonitrile solution) and heated to 37 ºC for 10 min with 1200 rpm shaking. The solution was then transferred to ultracentrifuge tubes (Beckman-Coulter 5/16” × 1 3/8”) and centrifuged at 53,000 x *g* for 20 min. Supernatant (30 μL) was carefully drawn, transferred to 0.2 mL amber TPX screw-cap 9 mm vials with short-thread conical glass inserts, sealed and stored at -20 ºC till LC-MS/MS analyses.

***P. putida* G7 chemoproteomic acquisition.** Peptide separation was executed on an EASY-nLC nanoflow ultra performance LC (UPLC) system coupled to an Orbitrap Fusion (Thermo Fisher Scientific) Tribrid MS. Analyses were performed in positive ion mode using the following LC-MS method: 8 μL injection volume, a positive ion spray voltage of 2500 V, static gas mode, an ion transfer tube temperature of 300 ºC. Peptides were injected into a reversed-phase C18 column (75 μm i.d. x 15 cm, 2 μm particles) and separated with a linear gradient of 0.1% formic acid in water (Buffer A) and 0.1% formic acid in 80% acetonitrile / 20% water (Buffer B). Chemoproteomic samples’ chromatographic elution was driven at a continuous flow rate of 330 nL/min via a multi-step linear gradient structured as follows: initialized at 2% Buffer B at 0 min, held at 2% for additional 2 min, stepped to 25% Buffer B over 50 min, resolved dynamically to 40% Buffer B over 10 min, and ramped to 95% Buffer B at 63 min where it was held static until 64 min to wash the analytical matrix.

The MS was operated in Data-Dependent Acquisition (DDA) mode. Survey scans (MS^1^) were obtained in the Orbitrap analyzer at a resolution of 120,000 (at 350 - 1400 *m/z*) with an automatic gain control (AGC) target of 8 x 10^5^ and a maximum injection time of 50 ms. Precursor selection for downstream tandem mass spectrometry (MS/MS) was executed across a 3 s cycle time. Precursor ions above 1 x 10^4^ intensity threshold were isolated utilizing a 1.6 *m/z* isolation window and subsequently fragmented via Higher-energy Collisional Dissociation (HCD) at a normalized collision energy (NCE) of 30%. Fragment spectra (MS^2^) were acquired in the Orbitrap at a resolution of 15,000 using a maximum injection time of 35 ms. Precursor ions were fragmented once, then set to a dynamic exclusion of 30 s, with an exclusion mass tolerance of ±5 ppm to minimize redundant sequencing.

**Chemoproteomic and metaproteomic profiling of gut microbial proteins from the small and large intestine.** Peptide separation was executed on a Vanquish Neo nano-UHPLC system coupled online to an Orbitrap Exploris 480 (Thermo Fisher Scientific) mass spectrometer. Analyses were done in positive ion mode using the following LC-MS method parameters: an 8 μL injection volume for chemoproteomic samples, a 4 μL injection volume for global metaproteomic samples, a positive ion spray voltage of 2500 V, and an ion transfer tube temperature of 280 ºC. Peptides were loaded and separated on a reversed-phase C18 analytical column (75 μm i.d. x 50 cm, 2 μm particles) using mobile phase consisting of 0.1% formic acid in water (Buffer A) and 0.1% formic acid in 80% acetonitrile / 20% water (v/v) (Buffer B). Global samples’ chromatographic elution was driven at a continuous flow rate of 300 nL/min via a multi-step linear gradient structured as follows: initialized at 1% Buffer B at 0 min, stepped to 3% Buffer B at 0.5 min, resolved dynamically to 36% Buffer B over 120 min, and ramped to 99% Buffer B at 121 min where it was held static until 130 min to wash the analytical matrix. Chemoproteomic samples’ chromatographic elution was driven at a continuous flow rate of 300 nL/min via a multi-step linear gradient structured as follows: initialized at 1% Buffer B at 0 min, stepped to 3% Buffer B at 0.5 min, resolved dynamically to 36% Buffer B over 60 min, and ramped to 99% Buffer B at 61 min where it was held static until 70 min to wash the analytical matrix.

The MS was operated in Data-Dependent Acquisition (DDA) mode. Survey scans (MS^1^) were obtained in the Orbitrap analyzer across a scan range of 375–1475 m/z at a resolution specification of 120,000, with an automatic gain control (AGC) target value of 3 x 10^6^. Precursor selection for downstream tandem mass spectrometry (MS/MS) was executed across a 3 s cycle time. Precursor ions exceeding an absolute intensity threshold of 5 x 10^3^ were isolated utilizing a 1.6 *m/z* isolation window and subsequently fragmented via Higher-energy Collisional Dissociation (HCD) at a normalized collision energy (NCE) of 30%. Fragment spectra (MS^2^) were recorded in the Orbitrap at a resolution capacity of 15,000 using an automated, auto-set maximum ion accumulation duration. Precursor ions were fragmented once, then set to a dynamic exclusion of 30 s enforcing an exclusion mass tolerance of ±10 ppm to minimize redundant sequencing.

**S3. Quantitative Metaproteomics and Chemoproteomics Data Analyses.**

Missing values were addressed within conditions using a hybrid strategy keyed to the probable mechanism of missingness, utilizing a fixed random seed of 1234. For the metaproteome datasets, values in a condition were assumed missing at random (MAR) and imputed via a $k$-nearest-neighbors ($k$NN) algorithm ($k=5$; row and column missingness caps of 0.50 and 0.80, respectively) when at most half of that condition’s replicates were missing. When more than half of the replicates within a condition were missing, or when a protein was completely absent from an entire experimental group, values were treated as missing not at random (MNAR) and imputed using a probabilistic minimum-detection estimator (MinDet, $q=0.05$). For the murine and single-organism chemoproteome datasets (four replicates per group), a matching hybrid framework was enforced: ($k$NN) ($k=5$) was utilized for proteins missing only one or two values in a group (MAR), while a probabilistic minimum-value estimator (MinProb, $q=0.01$) was applied for proteins missing three or four values in a group (MNAR), utilizing a group global-minimum fallback ($q=0.01$) for any residual missing values.

***Pseudomonas putida* G7 chemoproteomic analysis.** *P. putida* chemoproteomic intensities were characterized using the **CR-C** resin and the two naphthalene-derived resin probes (**Br-NAP** and **DADPS-Br-NAP**). Three limma contrasts were tested: Probe 1 (**BrNAP**) versus **CR-C**, Probe 2 (**DADPS-BrNAP**) versus **CR-C**, and Probe 1 versus Probe 2. The first two contrasts assessed protein enrichment by each probe relative to the control resin. The third directly compared protein recovery between the two probes. Proteins were considered probe-enriched relative to **CR-C** at an adjusted P value < 0.05 and a positive log2 fold change ≥ 1.5. For the direct probe comparison, an adjusted P value < 0.05 and an absolute log2 fold change ≥ 1.5 indicated preferential recovery by one probe, with the sign of the fold change indicating the favored probe.

**Multivariate and cross-omics analyses.** Exposure-associated structure was evaluated separately for the 16S rRNA gene and metaproteomic datasets by permutational multivariate analysis of variance using vegan::adonis2 with 9,999 permutations. Bray–Curtis dissimilarities were calculated from taxon relative abundances, whereas Euclidean distances were calculated from centered and standardized protein intensities. Homogeneity of multivariate dispersion was evaluated using betadisper and permutest with 9,999 permutations. Agreement between matched 16S rRNA gene and metaproteomic ordination structures was evaluated using Procrustes analysis with vegan::protest and by Mantel correlation between the corresponding distance matrices using Spearman’s rank correlation and 9,999 permutations.

**S4. Functional annotation and KO-level analysis.**

Each KO was assigned to one primary functional bin using KEGG pathway and BRITE information together with a predefined curated mapping (see Additional files 3 and 4). In parallel with the primary functional bins, ten mechanism-aware modules were curated to focus specific analyses on biologically relevant sets: central carbon metabolism, lipid metabolism, short-chain fatty acid and fermentation metabolism, xenobiotic and detoxification metabolism, aromatic amino acid metabolism, mucin and host-glycan degradation, bile acid biotransformation, redox metabolism, glutamate and GABA metabolism, and dietary glycan degradation. Complete KO assignments and targeted-module definitions are provided in Additional files 3 and 4.

KO abundance was calculated using non-imputed protein (NI) intensities. Linear protein intensities were summed across all proteins assigned to the same KO within each biological replicate. KO fold changes were calculated from the ratio of mean linear abundance in exposed and control mice. KOs detected in at least four of the eight replicates of either exposure group were compared using two-sided Wilcoxon rank-sum tests, with rank-biserial correlation reported as the effect size. *P* values were adjusted by the Benjamini–Hochberg procedure within each prespecified functional category. Where a unified, abundance-weighted response was required for a specific KO or a taxon-by-module combination, an abundance-aware differential response metric was computed as the abundance-weighted mean of the member proteins' per-protein $\log_{2}$ fold changes. For member proteins $i$ displaying a finite $\log_{2}$ fold change $L\left( i \right)$ and a strictly positive linear-scale abundance weight $w\left( i \right)$ (defined as the mean linear-scale abundance of protein $i$), the Abundance-Weighted Fold Change ($WFC$) was calculated as:

$$WFC=\frac{\sum_{i} w\left( i \right)\cdot L\left( i \right)}{\sum_{i} w\left( i \right)}$$

where the sums run over member proteins meeting these criteria, ensuring highly expressed proteins dominate the aggregate direction in proportion to measured linear abundance rather than log-scale values.

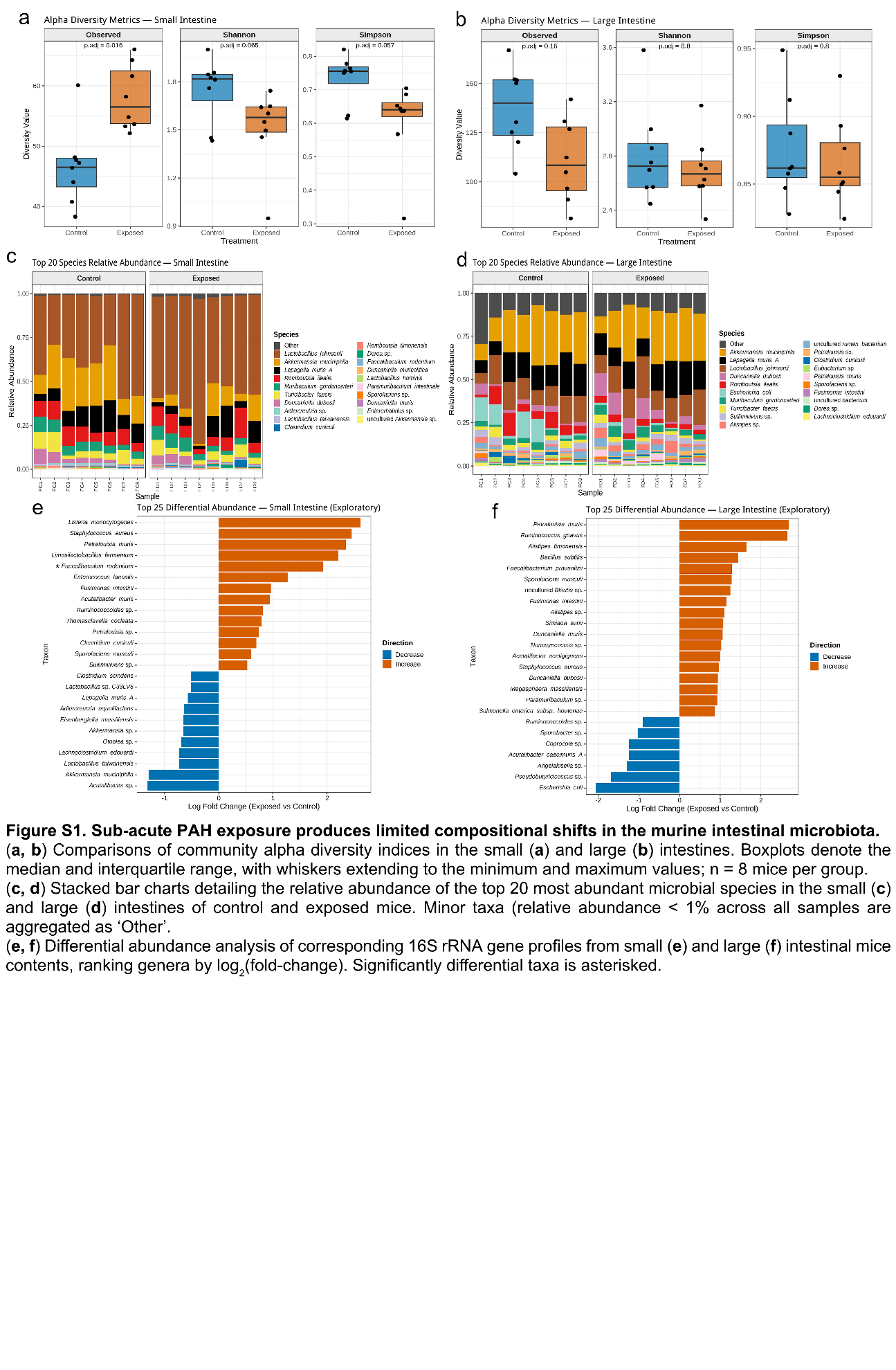

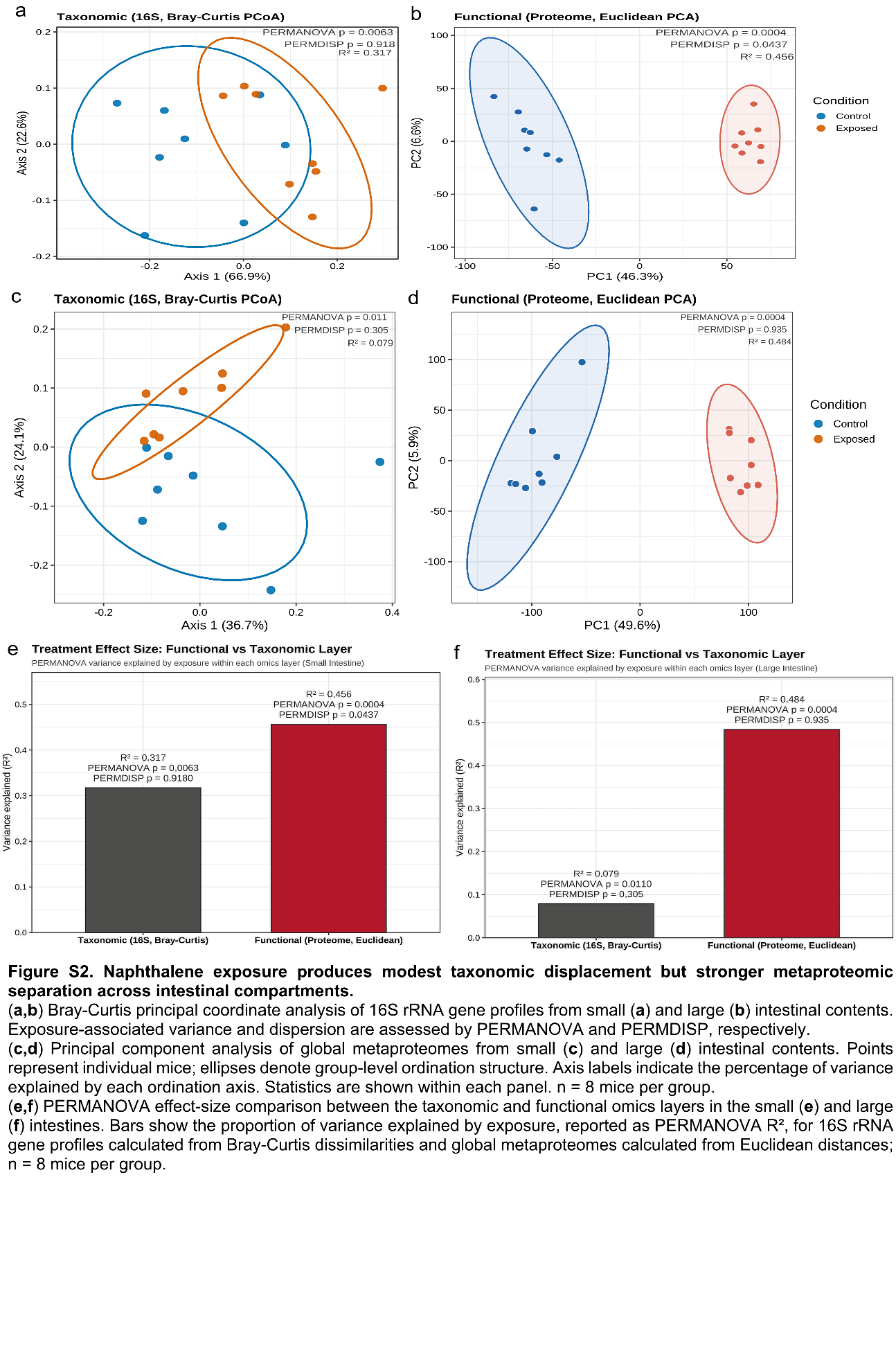

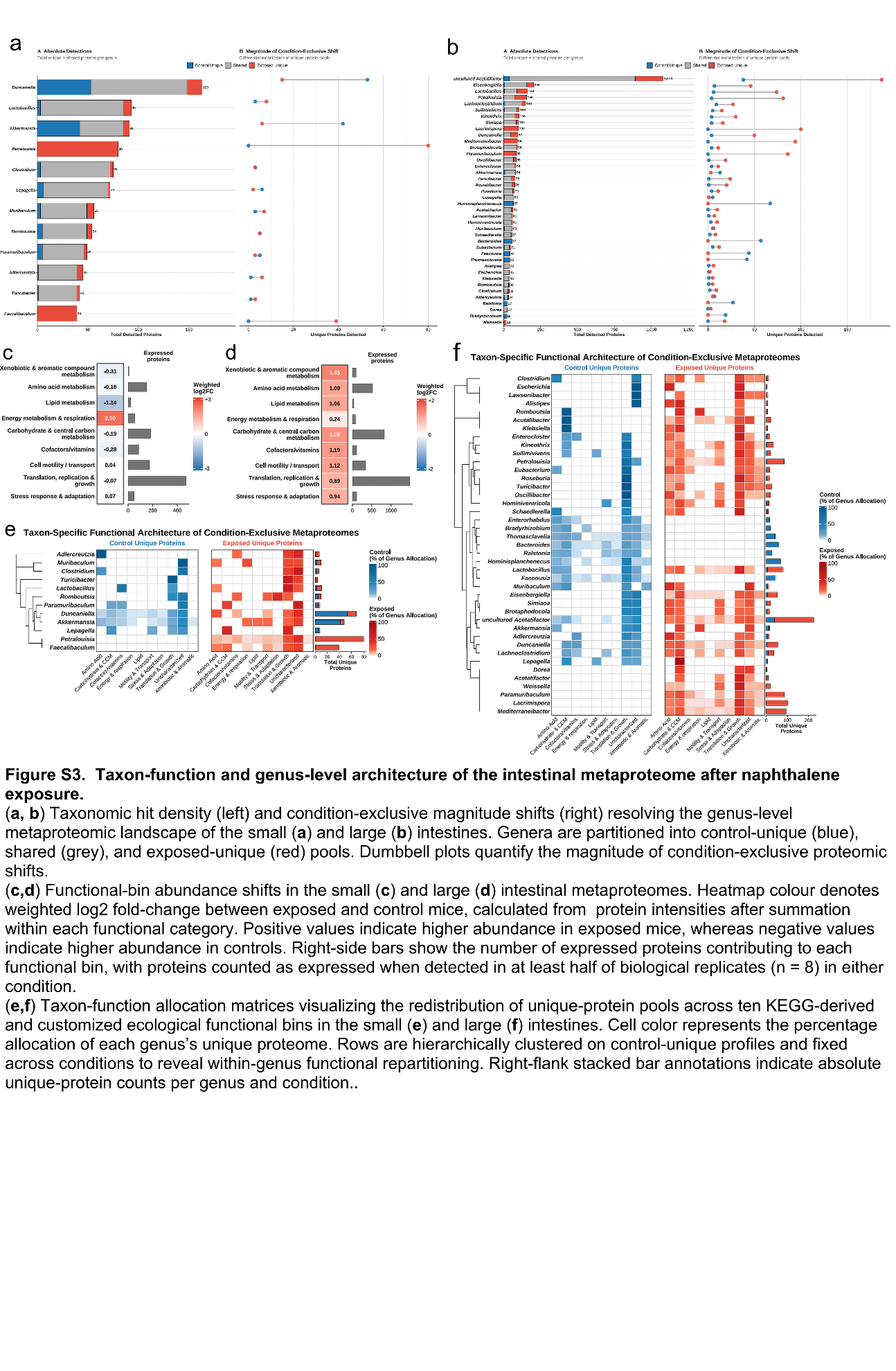

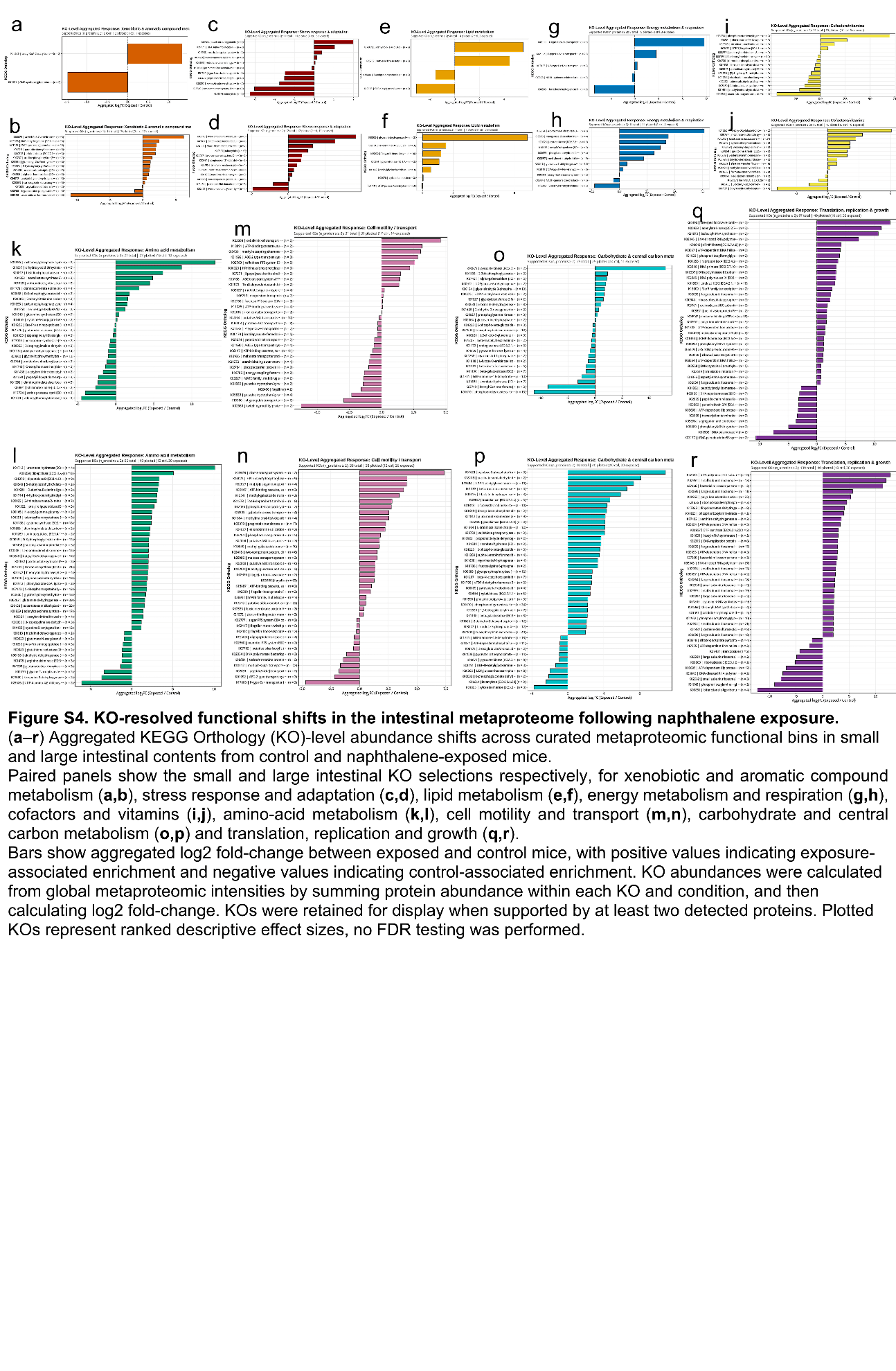

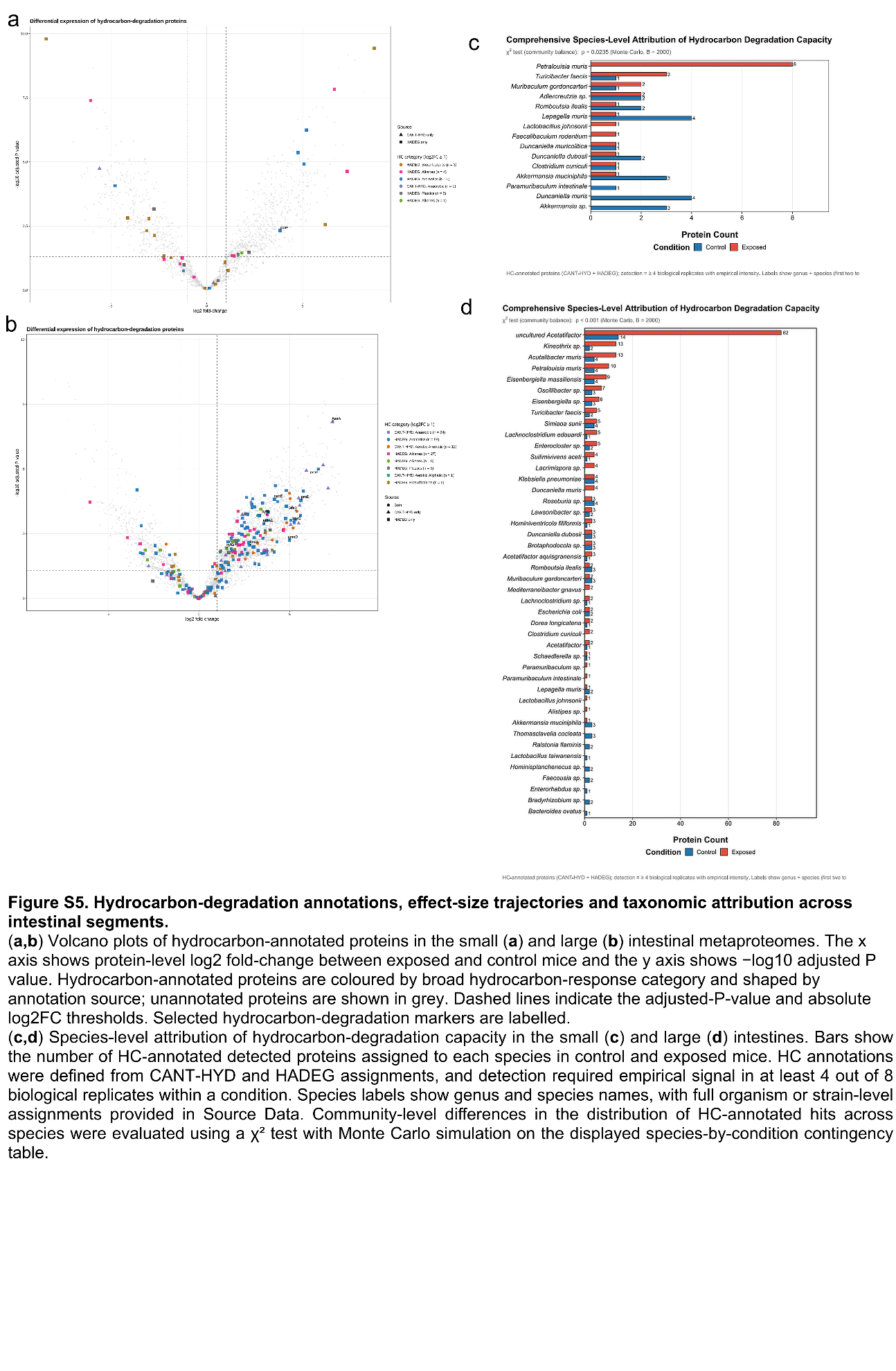

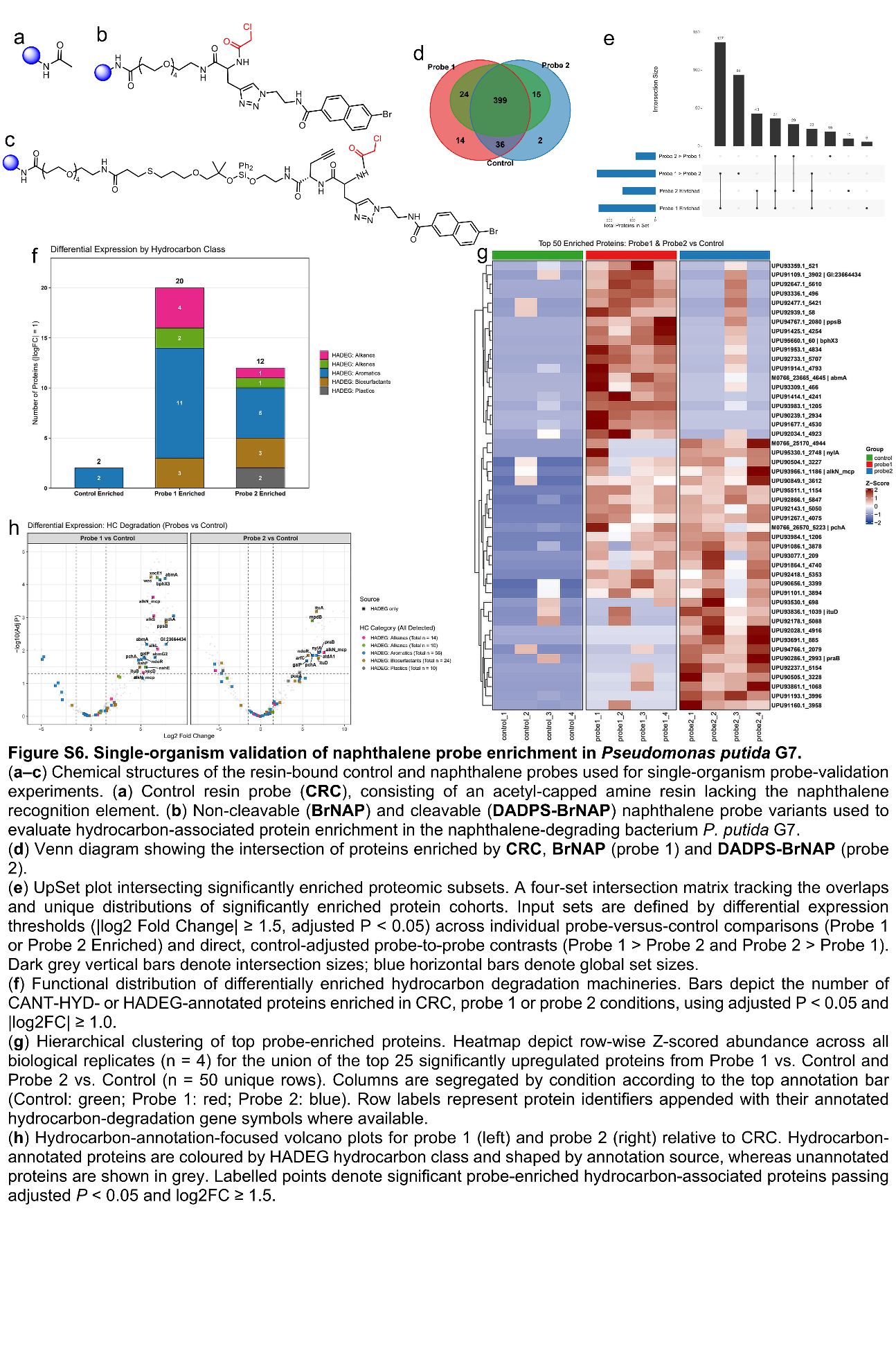

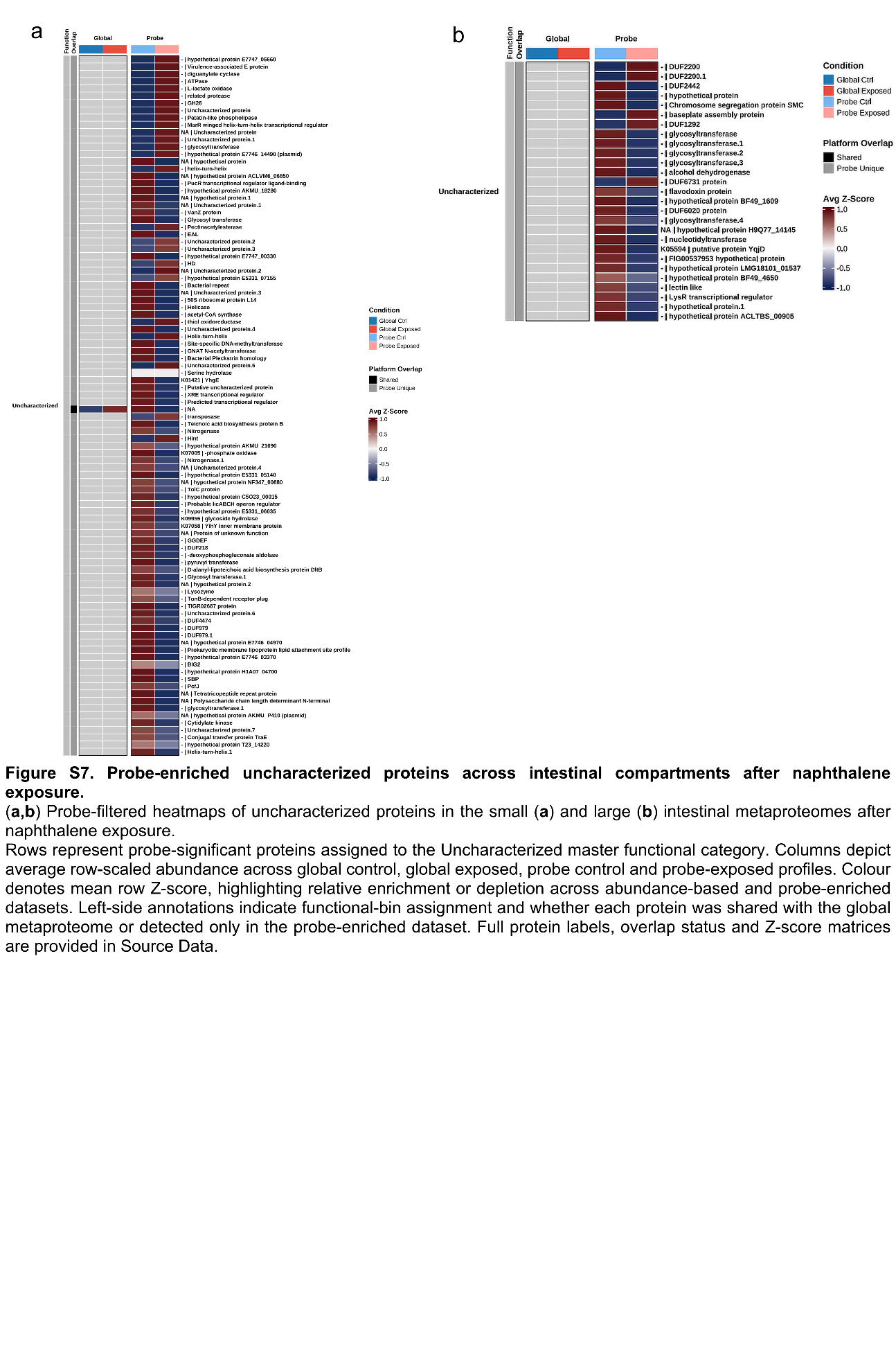

**S4. Synthetic Details**

**General Methods**

All solution-phase reactions were performed in oven-dried glassware under an atmosphere of dry N_2_ using standard Schlenk techniques unless otherwise stated. Anhydrous solvents were purchased in anhydrous grade and used as received. Reagents were purchased from commercial suppliers and used without further purification unless otherwise noted. Thin-layer chromatography (TLC) was performed on Supelco silica gel 60 F_254_ glass plates with UV (254 nm) or appropriate stains for visualization. Flash column chromatography was carried out on silica gel using the indicated solvent systems. ^1^H and ^13^C NMR spectra were collected in CDCl_3_ or DMSO-d_6_ (Sigma-Aldrich) on a Bruker 600 MHz spectrometer in the NMR facility, Baylor Sciences Building, Baylor University, Waco, Texas. Chemical shifts are reported in parts per million (ppm) relative to residual solvent signals of the deuterated solvents: CDCl_3_ (δ 7.26 for ^1^H, 77.16 for ^13^C) or DMSO-d_6_ (δ 2.50 for ^1^H, 39.52 for ^13^C). Splitting patterns are described as br (broad), s (singlet), d (doublet), t (triplet), q (quartet), m (multiplet), dd (doublet of doublets), and dt (doublet of triplets). Coupling constants (J) are reported in Hertz (Hz). High-resolution mass spectrometry was performed on a Thermo Orbitrap Q-Exactive using electrospray ionization (ESI). Reported m/z values correspond to the monoisotopic mass.

**Abbreviation List**

DCM	dichloromethane

DIPEA	*N,N*-diisopropylethylamine

DMF	*N,N*-Dimethylformamide

Et_3_N	triethylamine

Et_2_O	diethyl ether

EtOAc	ethyl acetate

Fmoc	fluorenylmethoxycarbonyl

HATU	hexafluorophosphate azabenzotriazole tetramethyluronium

NHS	*N*-Hydroxysuccinimde

PEG	polyethylene glycol

T3P	propanephosphonic acid anhydride

TFA	trifluoroacetic acid

THF	tetrahydrofuran

THPTA	Tris (3-hydroxypropyltriazolylmethyl)amine

**S5. Synthetic Schemes to Access Resin-immobilized Naphthalene Probes**
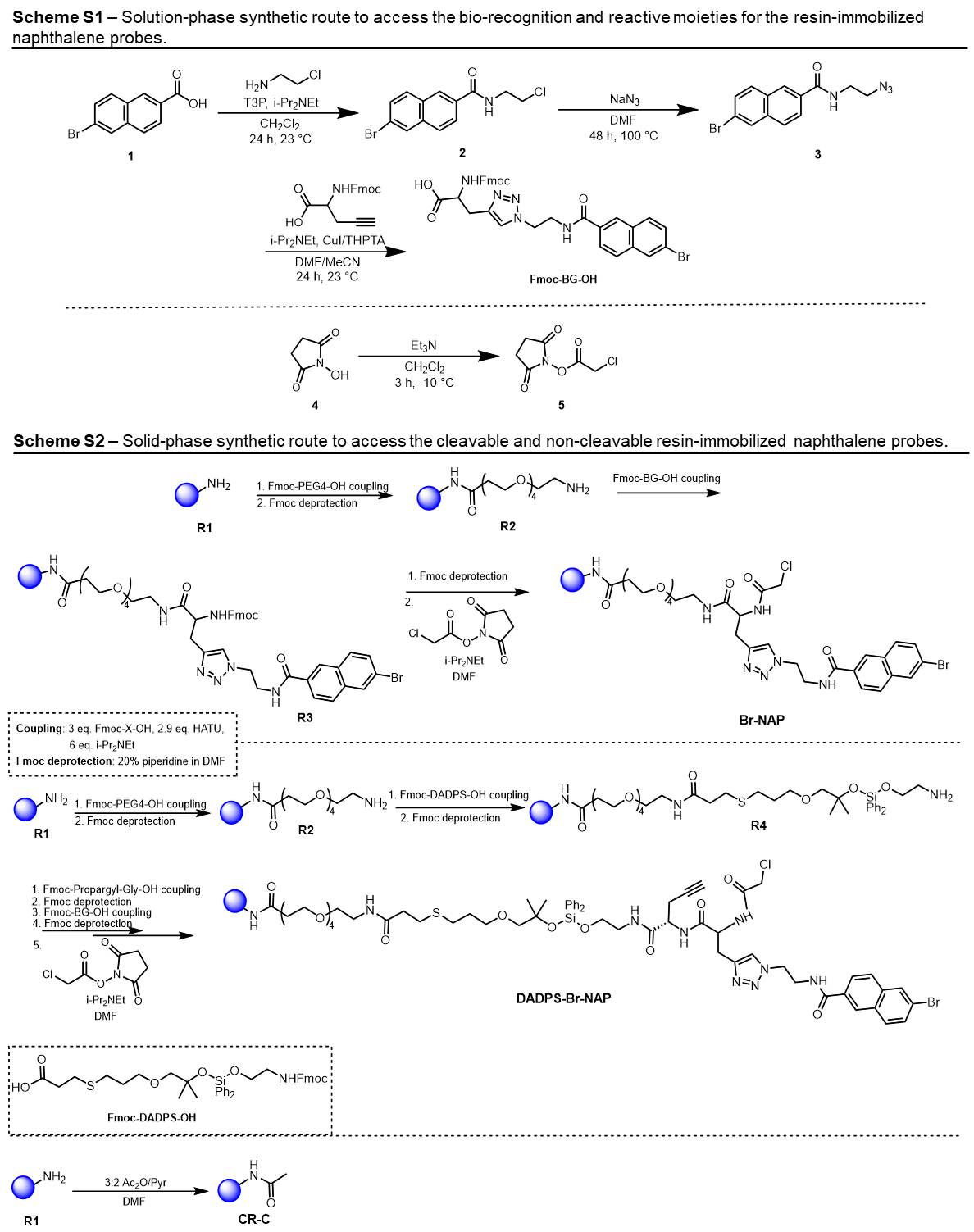

**Solution-Phase Synthetic Procedures and Characterization**

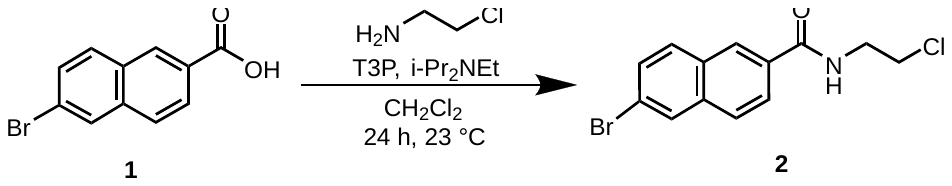

**Synthesis of Compound 2**

The synthesis followed an adapted procedure from[6].

*6-bromo-N-(2-chloroethyl)-2-naphthamide*.

6-Bromo-2-Naphthoic acid (**1**) (1.0 equiv) and 2-chloroethanamine (1.2 equiv) were weighed into an oven dried round bottomed flask equipped with a stir bar and capped. Vacuum purging and nitrogen backfilling were performed thrice. CH_2_Cl_2_ was added followed by DIPEA (2.0 equiv) and mixture stirred, followed by the addition of T3P (1.0 equiv) dropwise. Reaction stirred for 24 h with TLC monitoring till completion. Mixture was diluted with CH_2_Cl_2_ and washed with ice H_2_O, sat. NaHCO_3_ and brine solution. Organic layer was subsequently dried over sodium sulfate and concentrated *in vacuo*. Crude material was purified by flash column chromatography on silica with hexanes/ethyl acetate gradient, yielding the desired product as a white solid (1.01 g, 65%).

**^1^H NMR** (600 MHz, CDCl3) δ 8.28 (t, *J* = 1.3 Hz, 1H), 8.06 (d, *J* = 1.9 Hz, 1H), 7.87 (dd, *J* = 8.5, 1.7 Hz, 1H), 7.82 (t, *J* = 8.8 Hz, 2H), 7.63 (dd, *J* = 8.7, 1.9 Hz, 1H), 6.67 (s, 1H), 3.88 (td, *J* = 6.0, 4.9 Hz, 2H), 3.79 (dd, *J* = 6.0, 4.9 Hz, 2H).

**^13^C NMR** (151 MHz, CDCl_3_) δ 167.47, 135.90, 131.83, 131.15, 130.63, 130.52, 130.05, 127.80, 127.67, 124.73, 122.25, 41.95.

**HRMS (ESI-MS)** m/z: Calculated [M + Na]^+^ = 333.9605, Found [M + Na]^+^ = 333.9603

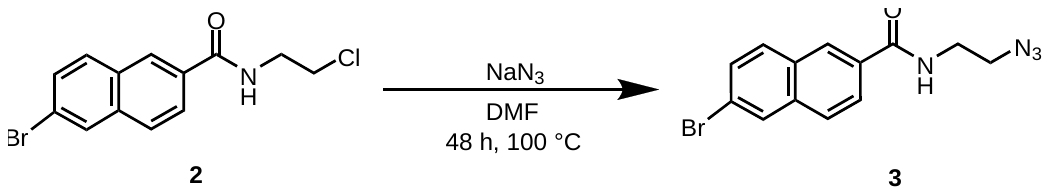

**Synthesis of Compound 3**

*N-(2-azidoethyl)-6-bromo-2-naphthamide*.

Compound **2** (1.0 equiv) and sodium azide (2.0 equiv) were weighed into an oven dried 2-neck round bottomed flask equipped with a stir bar. A reflux condenser fitted into one neck while the other was capped. Vacuum purging and nitrogen backfilling were performed thrice. DMF was added and solution heated to 100 ^o^C with stirring for 48 h. Upon reaction completion (monitored by TLC), the mixture was diluted with EtOAc and washed with 4% Aqueous Na_2_CO_3_ solution. Organic layer was dried over sodium sulfate, evaporated to isolate a white solid (446 mg, 85%) which was used in the next step without further purification.

**HRMS (ESI-MS)** m/z: Calculated [M + Na]^+^ = 341.0008, Found [M + Na]^+^ = 340.9992.

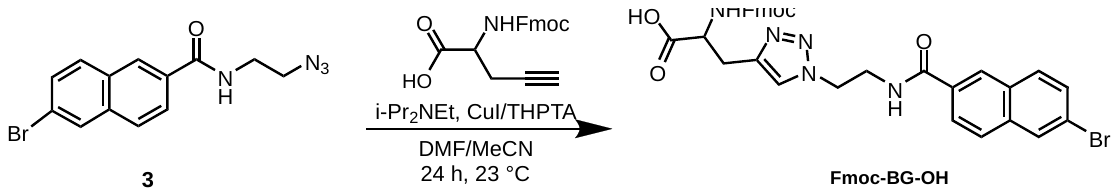

**Synthesis of Fmoc-BG-OH**

*2-((((9H-fluoren-9-yl)methoxy)carbonyl)amino)-3-(1-(2-(6-bromo-2-naphthamido)ethyl)-1H-1,2,3-triazol-4-yl)propanoic acid*.

Compound **3** (1.1. equiv), 2-((((9H-fluoren-9-yl)methoxy)carbonyl)amino)pent-4-ynoic acid (1.0 equiv), CuI (0.1 equiv), and THPTA (0.1 equiv) were weighed into an oven dried round bottomed flask equipped with a stir bar and capped. Vacuum purging and nitrogen backfilling were performed thrice. Anhydrous solution of DMF/MeCN (1:1 v/v) was added followed by DIPEA (2.0 equiv) and mixture stirred overnight. Upon completion, reaction mixture was concentrated *in vacuo*, then residue was diluted in ice H_2_O and extracted with CH_2_Cl_2_. Organic layer was washed with 1N HCl, sat. NH_4_Cl, and brine solution before being dried over sodium sulfate and concentrated. The crude material was purified by flash column chromatography on silica with CH_2_Cl_2_/MeOH gradient, yielding the desired product as a white solid (560 mg, 67%).

**^1^H NMR** (600 MHz, DMSO) δ 8.95 (t, *J* = 5.5 Hz, 1H), 8.47 (s, 1H), 8.23 (d, *J* = 2.1 Hz, 1H), 7.99 (d, *J* = 8.8 Hz, 1H), 7.95 (s, 2H), 7.86 (d, *J* = 7.6 Hz, 2H), 7.82 (s, 1H), 7.66 (dd, *J* = 8.7, 2.0 Hz, 1H), 7.62 (d, *J* = 7.6 Hz, 2H), 7.38 (t, *J* = 7.5 Hz, 2H), 7.28 (td, *J* = 7.3, 4.4 Hz, 2H), 6.96 (br s, 1H), 4.52 (t, *J* = 6.4 Hz, 2H), 4.19 – 4.13 (m, 3H), 4.03 (dd, *J* = 10.9, 5.7 Hz, 1H), 3.70 (t, *J* = 5.7 Hz, 2H), 3.17 – 3.11 (m, 1H), 2.99 (dd, *J* = 14.6, 7.6 Hz, 1H).

**^13^C NMR** (151 MHz, DMSO) δ 173.36, 166.41, 166.34, 155.48, 143.93, 143.83, 140.63, 135.23, 132.03, 131.10, 130.66, 129.69, 129.53, 128.92, 127.67, 127.56, 127.13, 127.06, 125.32, 125.23, 122.99, 121.39, 120.95, 120.06, 109.80, 65.44, 54.99, 48.35, 46.62, 28.34.

**HRMS (ESI-MS)** m/z: Calculated [M + Na]^+^ = 676.1166, Found [M + Na]^+^ = 676.1162

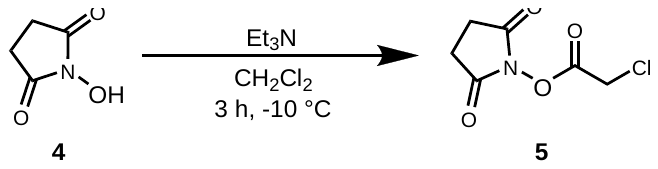

**Synthesis of Compound 5 (NHS-chloroacetyl ester)**

*Synthesis of 2,5-dioxopyrrolidin-1-yl 2-chloroacetate* (NHS-chloroacetyl ester) was carried out as previously described[7].

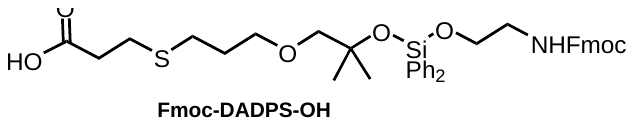

**Synthesis of Fmoc-DADPS-OH (Solid-Phase Compatible DADPS linker)**

*1-(9H-fluoren-9-yl)-10,10-dimethyl-3-oxo-8,8-diphenyl-2,7,9,12-tetraoxa-16-thia-4-aza-8-silanonadecan-19-oic acid* (Fmoc-DADPS-OH) linker was synthesized as previously described[8].

**Solid-Phase Organic Synthesis (SPOS) Protocol**

All solid-phase syntheses were carried out manually in either 10 mL fritted peptide vessels or polypropylene fritted Torviq syringes on TentaGel-XV-NH_2_ resin (theoretical loading 0.27 mmol/g). Resin beads were initially washed twice with DMF, then pre-swollen in DMF for 30 min before coupling reactions. Standard Fmoc SPPS protocol was used for coupling[5]. Couplings were done with 3 equiv of Fmoc-protected acid derivatives, 2.9 equiv of HATU, and 6 equiv of DIPEA. Kaiser tests were performed in between coupling or deprotection steps to ascertain completeness. Fmoc deprotections were carried out with 20% piperidine in DMF. A washing solution of DMF, MeOH, DCM, and DMF (3 x 1 min, with sufficient volume to fully cover the resin) was used.

**Resin loading determination**

The experimental loading of TentaGel-XV-NH_2_ resin (theoretical loading 0.27 mmol/g) was determined by UV-vis quantification of the Fmoc-piperidine adduct. Three samples of the resin (1.0 – 1.2 mg each) were transferred to glass vials and dried under a stream of nitrogen. Each sample was then treated with 1.5 mL of 20% piperidine in DMF (v/v) and allowed to stand at 23 ^o^C for 20 min, with occasional manual agitation. The resulting solutions were collected and their UV absorbance at 290 nm was recorded in triplicate using 20% piperidine in DMF as the blank.

The resin loading (mmol/g) was calculated as follows:

Fmoc loading (mmol/g) = A / (m x 1.75)

Where A is the measured absorbance at 290 nm and m is the mass of the resin in mg. The final loading value was obtained by taking the mean of the three measure samples. For the syntheses described herein, an experimental loading of 0.25 mmol/g was used.

**General coupling, deprotection, washing and capping protocol**

Prior to coupling, the resin was swelled in anhydrous DMF (ca. 10 mL per gram of resin) for 30 min at 23 ^o^C with gentle agitation. In a separate glass vial, the Fmoc-protected acid-functionalized compound (3.0 equiv relative to the resin loading) and HATU (2.9 equiv) were dissolved in DMF. DIPEA (6.0 equiv) was added, the mixture was stirred for 30 min to activate the acid before being added to the pre-swollen resin. The suspension was gently agitated at 23 ^o^C as coupling proceeded, with Kaiser test monitoring (as described below) to determine completion. Post coupling, the resin was drained and washed sequentially with the washing solution.

**General Fmoc deprotection**

Deprotection of Fmoc was performed with 20% piperidine in DMF (v/v) solution. The resin was first treated with the solution for 20 min at 23 ^o^C with gentle agitation, drained, and then treated with a fresh portion of the 20% piperidine in DMF for an additional 15 min (2x). Following the confirmation of deprotection via a Kaiser test, the resin was washed with the sequential washes as earlier described.

**Kaiser test for monitoring coupling and deprotection**

The Kaiser test (ninhydrin/phenol/KCN) was applied to qualitatively determine the completion of coupling and deprotection steps. For each test, a small portion of the resin beads was transferred to a glass vial and treated with 3 drops each of the Kaiser solutions A (ninhydrin), B (phenol) and C (KCN), then heated at 100 ^o^C for 5 min. Colorless or pale yellow beads indicated successful coupling while deep blue beads indicated successful deprotection.

**General capping of unreacted amines**

After each coupling, unreacted amines were capped by treating the resin with a solution of Ac_2_O/Pyr (3:2, v/v) in DMF for 30 min at 23 ^o^C with gentle agitation. The resin was then washed with the washing solution before proceeding with the next coupling.

**Preparation of probe-functionalized and control resins**

For **BrNAP** probe, sequential couplings of PEG spacer (Fmoc-N-amido-PEG_4_-OH) and Fmoc-BG-OH were performed to generate the resin-probe scaffold. Next, Fmoc deprotection was carried out and the NHS-chloroacetyl ester (compound **5**) was added with 2 equiv of DIPEA in DMF. The mixture was rotated for 2 h, then the resin was filtered, washed three times with DMF, DCM, MeOH, THF, and Et_2_O and dried in a desiccator. Dried resin was stored at 4^o^C before use.

For **DADPS-BrNAP** probe, sequential couplings of PEG spacer (Fmoc-N-amido-PEG4-OH), Fmoc-DADPS-OH, and Fmoc-N-amido-propargylglycine were performed to generate the resin-probe scaffold. Following Fmoc deprotection, Fmoc-BG-OH was coupled with corresponding Fmoc deprotection prior to the treatment of the resin with NHS-chloroacetyl ester (compound **5**) and 2 equiv of DIPEA in DMF. Subsequently, the resin was washed, dried and stored as described above.

For capped resin control (**CR-C**), TentaGel-XV-NH_2_ resin was acetylated with 3:2 equiv of Ac_2_O: Pyridine solution in DMF. Resin was filtered, washed three times with DMF, DCM, MeOH, THF, and Et_2_O and dried in a desiccator. Dried resin was stored at 4^o^C.

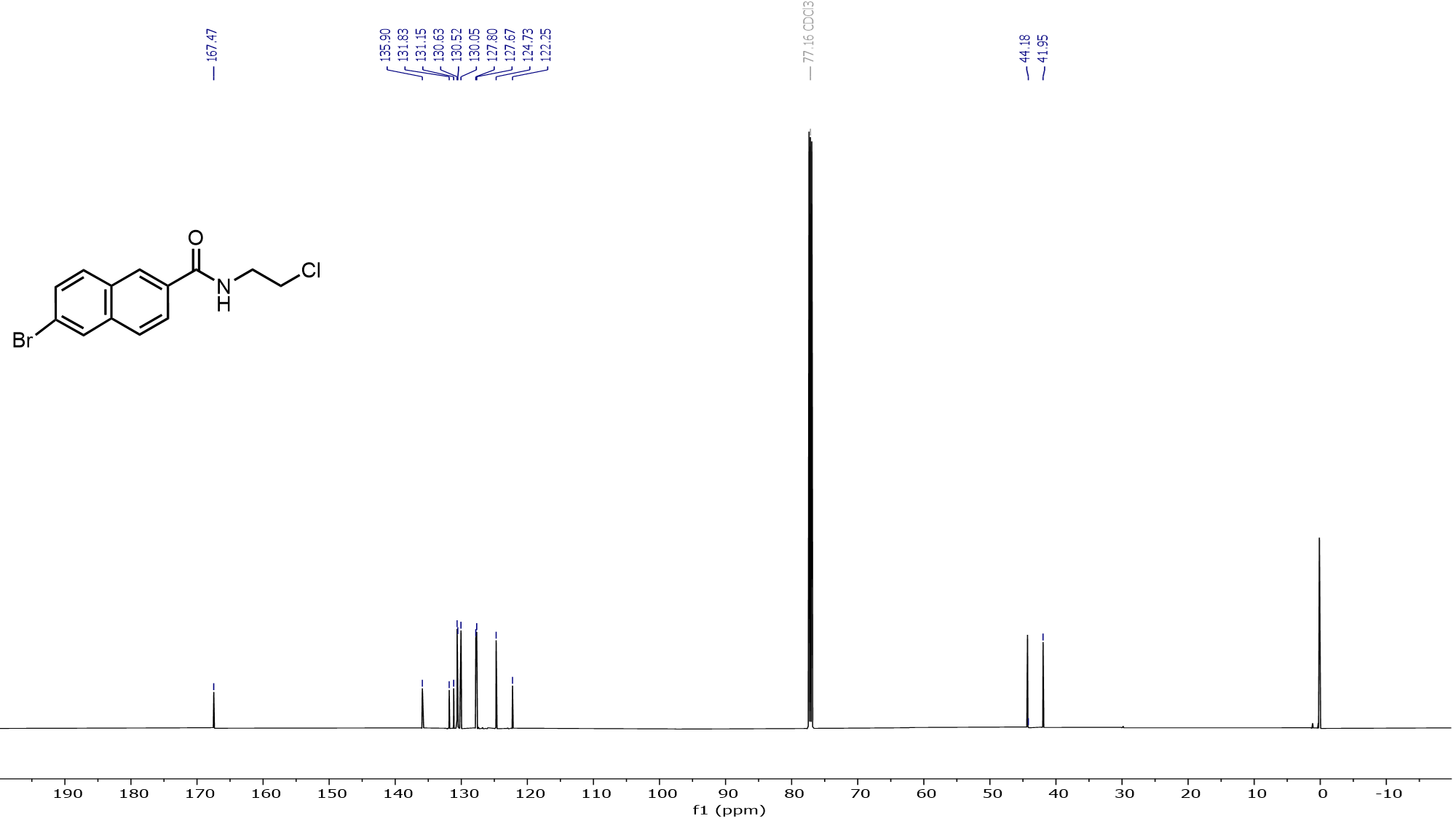

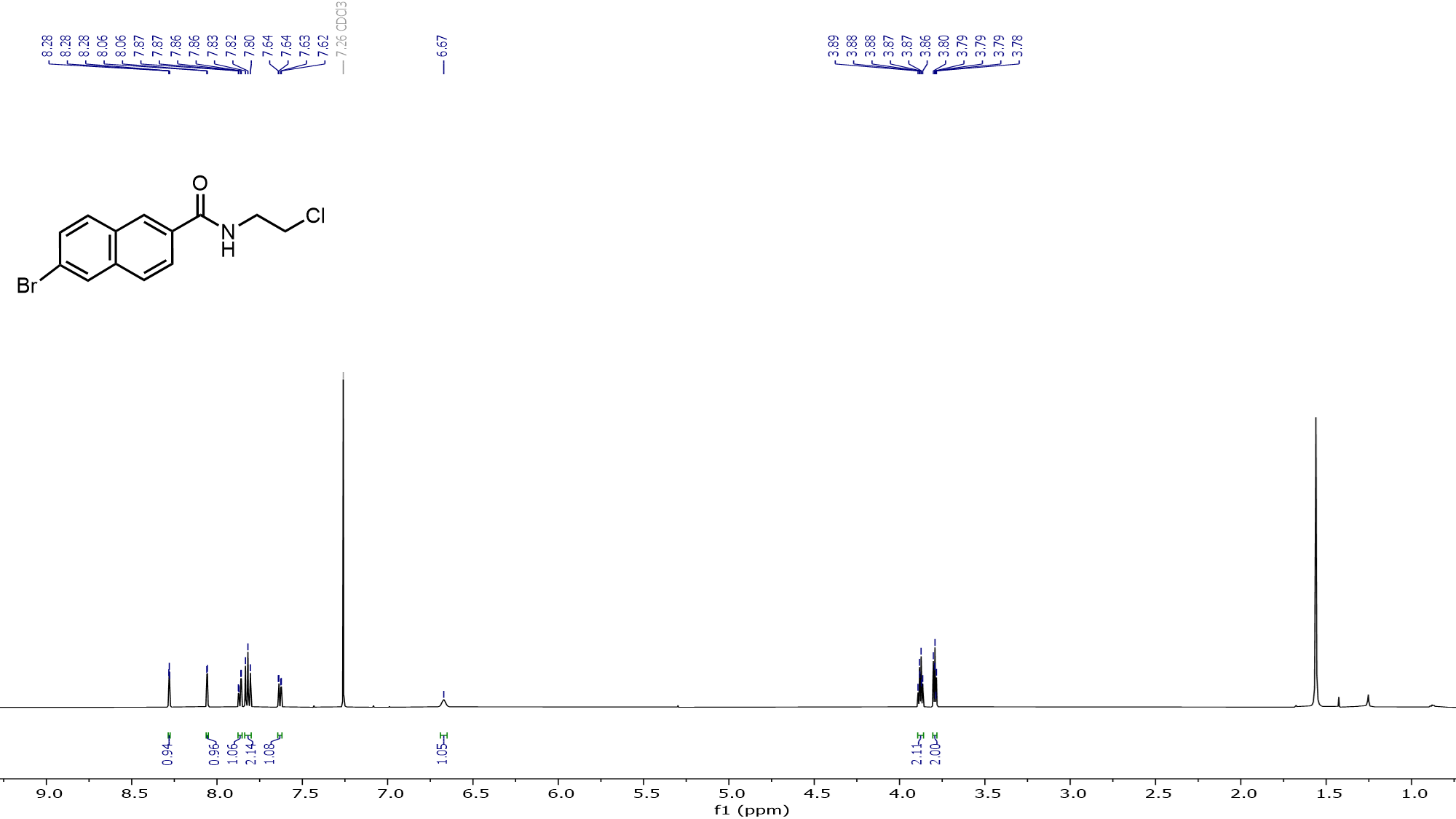

**S6. NMR Spectra**

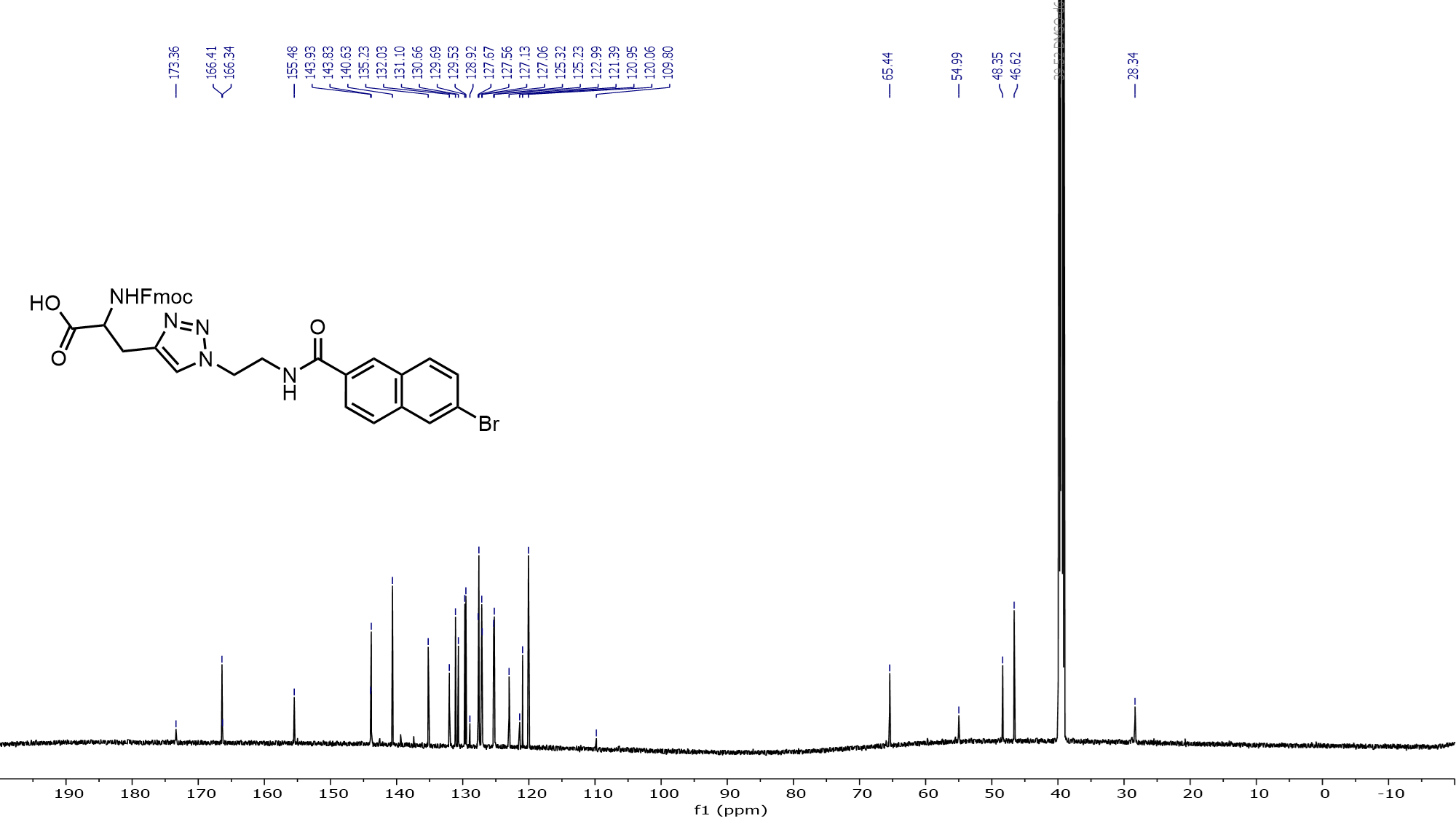

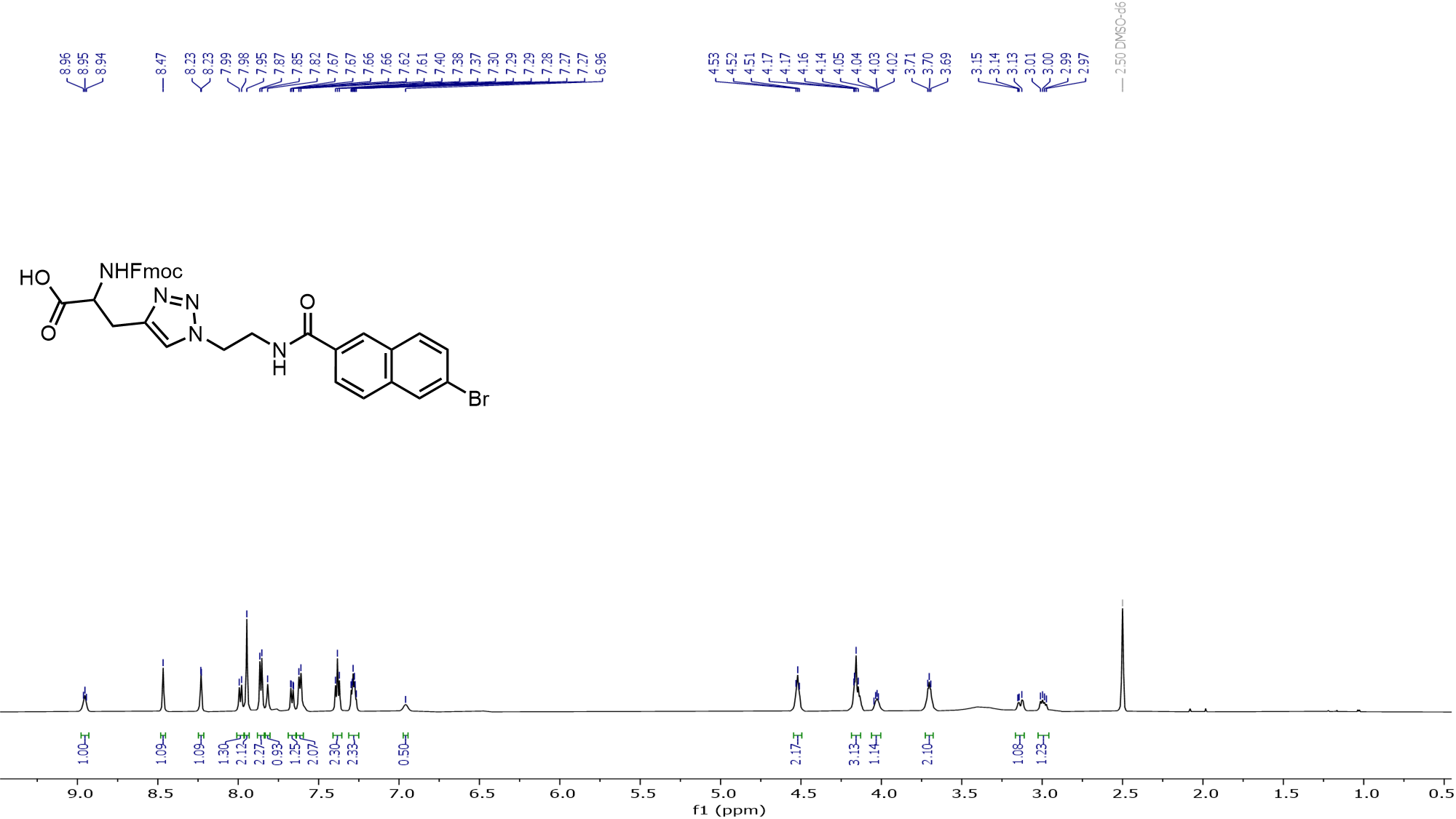
